# RAVEN: Robust, generalizable, multi-resolution structural MRI upsampling using Autoencoders

**DOI:** 10.1101/2025.09.22.677945

**Authors:** Walter Adame Gonzalez, Roqaie Moqadam, Yashar Zeighami, Mahsa Dadar

## Abstract

Due to their high inter-tissue contrast, Magnetic resonance images (MRIs) can reflect neuroanatomical changes related to healthy aging and pathological processes. However, standard brain MRI acquisition resolutions hinder the ability to measure the more subtle changes that occur in early disease stages. Increasing the resolution during acquisition poses multiple challenges, including increased noise, higher acquisition times and cost, and discomfort of the scanned individual. In this work, we propose a robust, generalizable single-image super-resolution network for brain MRIs named Resolution Augmentation with Variational auto-Encoder Networks (RAVEN) with generative adversarial networks (GANs). We show RAVEN is capable of upsampling in-vivo and ex-vivo MRIs of diverse modalities (e.g. T1-weighted, T2-weighted, and T2*) and varying field strengths (3T to 7T) to target voxel sizes as small as 0.5mm isotropic using arbitrary upsampling factors. RAVEN achieved state-of-the-art performance against deep learning and non-deep learning methods, best preserving true anatomical information. We have also made RAVEN open access, with the source code as well as training and evaluation scripts available and ready to use at: https://github.com/waadgo/raven.

## 1. Introduction

Magnetic Resonance Imaging (MRI) is a non-invasive technique widely used in clinical and research protocols for studying human brains in-vivo. In research settings, MRI scans are commonly acquired at voxel sizes of approximately 1mm isotropic and analyzed with automatic image processing pipelines (Dadar et al., n.d.; Iglesias et al., 2021; Lüsebrink et al., 2018). However, early detection of fine brain atrophy patterns requires accurate morphometry of small brain regions for which standard resolutions are suboptimal (Shafiee et al., 2024).

Acquisition voxel size is a hindering factor when performing brain morphometry, since it limits the minimum size of brain regions that would be measurable on an MRI. Therefore, it is desirable to acquire MRIs with the highest resolution possible. However, increasing acquisition resolution substantially increases noise level as well as time, noise, discomfort of scanned individuals, and makes the scan more prone to movement artifacts (Zhu et al., 2009). Therefore, in clinical studies and practical applications, it is uncommon to use high-resolution full brain MRI acquisitions. Furthermore, numerous legacy datasets have been scanned using standard-resolution sequences, but contain a great amount of valuable information. As an example. the the Alzheimer’s Disease Neuroimaging Initiative (ADNI) (Mueller et al., 2005), one of the largest databases of individuals on the continuum of Alzheimer’s Disease containing longitudinal demographic, genetic, and clinical information, as well as positron-emission tomography (PET) scans, provides longitudinal MRI data acquired at ∼1mm isotropic voxel sizes.

Single-Image Super-Resolution (SISR) is a technique to artificially increase the resolution of images after their acquisition, generating a high-resolution (HR) image from a single low-resolution (LR) input. SISR has gained popularity in naturalistic image processing (Fischer et al., 2024; Jo et al., 2020; Rombach et al., 2022; Shi et al., 2016; Umer et al., 2020; Umer & Micheloni, 2020; X. Wang et al., 2018; Zeng et al., 2017; Zhang et al., 2021), as well as in medical image processing (Avants et al., 2023; Y. Chen et al., 2018; Iglesias et al., 2021; Lin et al., 2023; J. Wang et al., 2023; K. Zhao et al., 2025). The latter implies potential valuable gains in accuracy for medical diagnostic and prognostic purposes, and could even make it possible to measure anatomical structures that would be too small to capture in the LR regime (Shafiee et al., 2024)}. SISR is an ill-posed problem since a single LR sample can be upsampled into many different HR images. Therefore, caution should be taken when utilizing image-reconstruction techniques in the medical field, and thorough validations should be performed before publicly releasing a SISR method.

Several strategies have been proposed for MRI SISR in the literature. Early non-learning approaches include Non-local Means Upsampling (NLMUPSAMPLE) (Manjón et al., 2010), a contrast- and resolution-agnostic self-similarity method that iteratively refines a 3D estimate while enforcing consistency (i.e., voxel-level similarity) between the input LR image and a downsampled version of the provisional HR image. More recently, deep learning (DL) has achieved state-of-the-art performance owing to its capacity to model non-linear mappings. For example, Chen and colleagues (Y. Chen et al., 2018) proposed a 3D densely connected convolutional neural network (CNN) for SISR of T1w MRIs using an upsampling factor of 2×2×2, demonstrating improvements over 2D CNN counterparts and interpolation. Another relevant study (Lyu et al., 2018) introduced a Generative Adversarial Network (GAN) trained on 2D patches for upsampling single-modality T2-weighted (T2w) MRIs, likewise outperforming interpolation. Additionally, Avants and colleagues (Avants et al., 2023) proposed ANTsPyNet for MRI SISR based on the previous work of Tustison and colleagues (Tustison et al., 2021) based on a 3D Deep Back-Projection Networks (DBPN) (Haris et al., 2018) for upsampling of T1-weighted (T1w) MRIs. More recently, Wang and colleagues (J. Wang et al., 2023) proposed brain MRI SISR using latent diffusion models, and later Zhao and colleagues (K. Zhao et al., 2025) proposed partial diffusion models, in which a compressed representation of the low-resolution input is degraded with normal gaussian noise and then gradually denoised to match the latent representation of a high-resolution MRI. While most other works for MRI SISR aimed for through-plane upsampling of 2D MRI samples (Lin et al., 2023; K. Zhao et al., 2025), or propose independent weights for a finite set of upsampling factors (Avants et al., 2023; Haris et al., 2018; J. Wang et al., 2023), Wu and colleagues (Wu et al., 2023) proposed ArSSR, an arbitrary scale super-resolution method for T1w brain MRIs capable of handling arbitrary upsampling factors for all three anatomical axes using an encoder-decoder approach.

Despite these advances, these prior works have been either focused on a single imaging modality (mainly T1w), and limited to specific resolutions and upsampling factors. As such, there still remains a gap in developing robust methods that are simultaneously modality, resolution, and upsampling-factor agnostic. The present work aimed to address these gaps. To achieve this, we trained a single network that is capable of processing multiple MRI contrasts, magnetic field strengths, in-vivo and ex-vivo images of in situ brains as well as ex situ hemispheres, enabling generalizability of the network to multiple inter-tissue contrasts. Additionally, we trained and validated our network using multiple input and target voxel sizes, modelling different degradation functions, and further validating on out-of-distribution data, ensuring the robustness and generalizability of its performance. Finally, we evaluated segmentation performance on the upsampled images as a downstream task after super-resolution, demonstrating the ability of our method to reconstruct true anatomical details.

## 2. Methods

### 2.1. Datasets

To train deep neural networks for brain MRI super-resolution that generalize across varying tissue contrasts, voxel sizes, and acquisition parameters (i.e., different modalities), it is essential to use datasets that are as diverse as possible. As reported in Table 1, we used an aggregate training dataset that includes native images acquired at isotropic voxel sizes ranging from 0.4mm to 1mm from in-vivo T1w, T2w, T2*, and T1-maps, and ex-vivo T1w and T2w head and hemisphere scans acquired at multiple resolutions (See Supplementary materials-Data Description section for a detailed description of the used datasets).

**Table 1.** Datasets used for training, validating, and testing RAVEN.

| Dataset | Split | Modality | Field strength | N | Voxel size [mm] | Individuals | Ages |
| --- | --- | --- | --- | --- | --- | --- | --- |
| AHEAD(Alkemade et al., 2020) | Train | T1 map | 7T | 66 | 0.7×0.7×0.7 | 66 | 18-80 |
|  |  | T1w | 7T | 66 | 0.7×0.7×0.7 |  |  |
|  | Validation | T1 map | 7T | 19 | 0.7×0.7×0.7 | 19 |  |
|  |  | T1w | 7T | 19 | 0.7×0.7×0.7 |  |  |
|  | Test | T1 map | 7T | 10 | 0.7×0.7×0.7 | 10 |  |
|  |  | T1w | 7T | 10 | 0.7×0.7×0.7 |  |  |
| AOMICID1000(Snoek et al., 2021) | Train | T1w | 3T | 648 | 1.0×1.0×1.0 | 648 | 19-26 |
|  | Validation | T1w | 3T | 185 | 1.0×1.0×1.0 | 185 |  |
|  | Test | T1w | 3T | 94 | 1.0×1.0×1.0 | 94 |  |
| AOMICPIOP1(Snoek et al., 2021) | Train | T1w | 3T | 151 | 1.0×1.0×1.0 | 151 | 19-26 |
|  | Validation | T1w | 3T | 43 | 1.0×1.0×1.0 | 43 |  |
|  | Test | T1w | 3T | 22 | 1.0×1.0×1.0 | 22 |  |
| AOMICPIOP2(Snoek et al., 2021) | Train | T1w | 3T | 158 | 1.0×1.0×1.0 | 158 | 19-26 |
|  | Validation | T1w | 3T | 45 | 1.0×1.0×1.0 | 45 |  |
|  | Test | T1w | 3T | 23 | 1.0×1.0×1.0 | 23 |  |
| CHEN(X. Chen et al., 2023) | Train | T1w | 3T | 7 | 0.8×0.8×0.8 | 7 | 25-41 |
|  |  | T1w | 7T | 7 | 0.65×0.65×0.65 |  |  |
|  | Validation | T1w | 3T | 2 | 0.8×0.8×0.8 | 2 |  |
|  |  | T1w | 7T | 2 | 0.65×0.65×0.65 |  |  |
|  | Test | T1w | 3T | 1 | 0.8×0.8×0.8 | 1 |  |
|  |  | T1w | 7T | 1 | 0.65×0.65×0.65 |  |  |
| DH DATASET<br>(post-mortem)(Dadar et al., 2024) | Train | T1w | 3T | 152 | varying | 103 | N/A |
|  |  | T2w | 3T | 152 | varying |  |  |
|  | Validation | T1w | 3T | 43 | varying | 34 |  |
|  |  | T2w | 3T | 43 | varying |  |  |
|  | Test | T1w | 3T | 23 | varying | 22 |  |
|  |  | T2w | 3T | 23 | varying |  |  |
| HBA(Schira et al., 2023) | Train | T1w | 7T | 1 | 0.5×0.5×0.5 | 1 | 30, 45 |
|  |  | T2w | 7T | 1 | 0.4×0.4×0.4 |  |  |
|  | Validation | T1w | 7T | 0 | 0.5×0.5×0.5 | 0 |  |
|  |  | T2w | 7T | 0 | 0.4×0.4×0.4 |  |  |
|  | Test | T1w | 7T | 1 | 0.5×0.5×0.5 | 1 |  |
|  |  | T2w | 7T | 1 | 0.4×0.4×0.4 |  |  |
| HCP(Van Essen et al., 2012) | Train | T1w | 3T | 773 | 0.7×0.7×0.7 | 773 | 22, 35 |
|  |  | T2w | 3T | 771 | 0.7×0.7×0.7 |  |  |
|  | Validation | T1w | 3T | 219 | 0.7×0.7×0.7 | 221 |  |
|  |  | T2w | 3T | 220 | 0.7×0.7×0.7 |  |  |
|  | Test | T1w | 3T | 219 | 0.7×0.7×0.7 | 221 |  |
|  |  | T2w | 3T | 220 | 0.7×0.7×0.7 |  |  |
| OSCBS(Tardif et al., 2016) | Train | T1w | 7T | 18 | 0.5×0.5×0.5 | 18 | 26-27 |
|  |  | T2* | 7T | 6 | 0.5×0.5×0.5 |  |  |
|  | Validation | T1w | 7T | 5 | 0.5×0.5×0.5 | 5 |  |
|  |  | T2* | 7T | 3 | 0.5×0.5×0.5 |  |  |
|  | Test | T1w | 7T | 3 | 0.5×0.5×0.5 | 3 |  |
|  |  | T2* | 7T | 0 | 0.5×0.5×0.5 |  |  |
| LUSENBRINK(Lüsebrink et al., 2018) | Train | T1w | 7T | 0 | 1.0×1.0×1.0 | 0 | 35 |
|  |  | T1w | 7T | 0 | 0.5×0.5×0.5 |  |  |
|  |  | T1w | 7T | 0 | 0.25×0.25×0.25 |  |  |
|  | Validation | T1w | 7T | 0 | 1.0×1.0×1.0 | 0 |  |
|  |  | T1w | 7T | 0 | 0.5×0.5×0.5 |  |  |
|  |  | T1w | 7T | 0 | 0.25×0.25×0.25 |  |  |
|  | Test | T1w | 7T | 1 | 1.0×1.0×1.0 | 1 |  |
|  |  | T1w | 7T | 1 | 0.5×0.5×0.5 |  |  |
|  |  | T1w | 7T | 1 | 0.25×0.25×0.25 |  |  |

### 2.2. Data preprocessing

Preprocessing consisted of denoising the images with FONDUE (Adame-Gonzalez et al., 2023), a DL-based method capable of denoising multiple input modalities and resolutions, followed by reorientation to Right-Anterior-Superior (RAS) and robust intensity rescaling between 0 and 255 (cropping the intensities beyond the 99.5 percentiles). The axes were then randomly swapped (changing the effective brain orientation) and a foreground mask was created through an intensity threshold.

Non-overlapping patches of shape 192×192×32 (center cropped during training to 160×160×32) were used for memory efficiency. The patch was centered in the high-resolution plane, with the sliding window for extracting the patches moving along the third axis. As expected, some patches contained no foreground information. Since the goal was to train a model that upsamples the foreground but still has some training on how to handle background patches, 50% of the patches containing less than 20% voxels with foreground information were randomly dropped. Finally, the individual patches were normalized to the intensity range of [-1, 1].

### 2.3. Architecture

We trained a Variational AutoEncoder Generative Adversarial Network (VAE-GAN) with small KL-regularization adapted from the method by Rombach and colleagues (Rombach et al., 2022) named <u>R</u>esolution <u>A</u>ugmentation with <u>V</u>ariational auto-<u>E</u>ncoder <u>N</u>etworks with GANs (RAVEN). Figure 1 shows the architecture of RAVEN.

**Figure 1.**
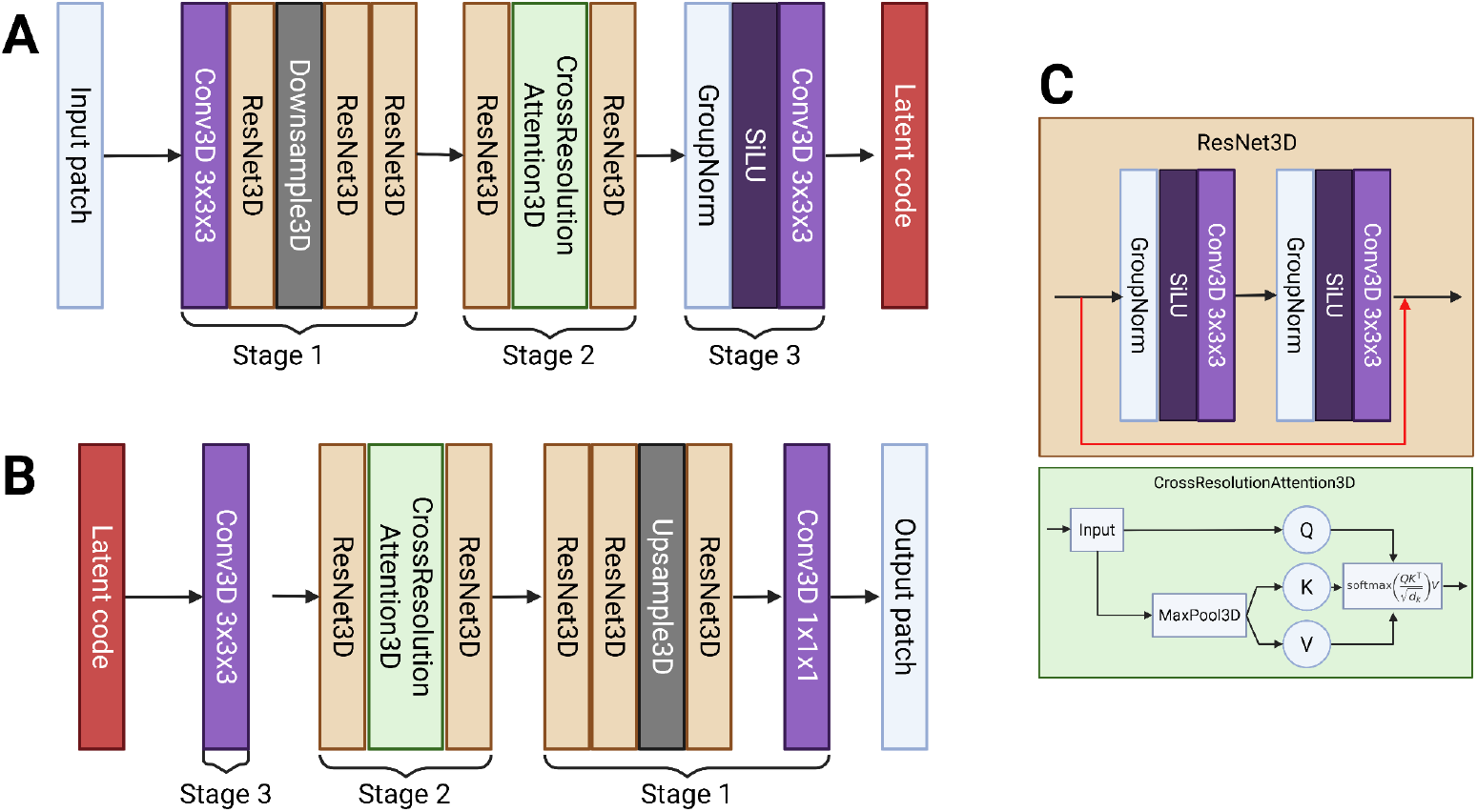
RAVEN architecture: Panel A depicts the encoder, receiving the input patch of dimensions (*D*) × (*H*) × (*W*) with three main stages and as output a 3D latent code of shape (*D* ÷ 2) × (*H* ÷ 2) × (*W* ÷ 2). Panel B shows the decoder receiving as input the latent code, and generating an output patch with dimensions (*D*) × (*H*) × (*W*). Panel C depicts the internal architecture of ResNet3D and the CrossResolutionAttention3D module.

**Figure 2.**
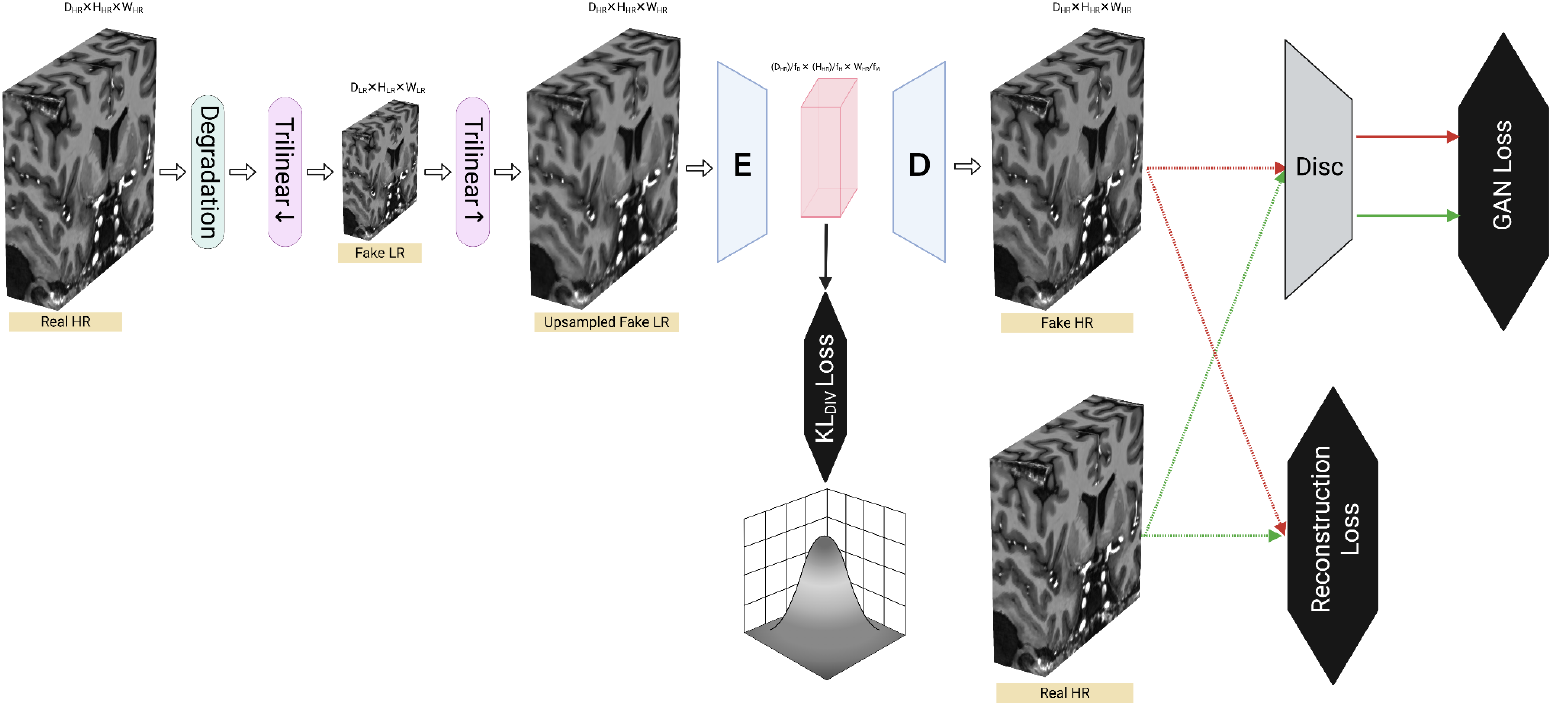
Training procedure for RAVEN: A surrogate of low-resolution image patch (Fake LR) is generated by degrading and downsampling (through trilinear interpolation) an input non-degraded image patch (Real HR). Then, the input to the encoder (E) is generated by resampling the Fake LR image to the target space using trilinear interpolation (Upsampled Fake LR). The latent space is a compressed representation of the input patch and is regularized using KL Divergence. The generated Fake HR and the Real HR patches get parsed to the discriminator network (Disc) to compute the discriminator loss and the GAN loss, as well as the Reconstruction loss.

### 2.3.1. Encoder

**The generator encoder** *G*_*E*_ consists of three stages (Figure 1-A):

<u>***Stage 1:***</u> Conv3D with 3×3×3 kernel, stride of 1 and padding of 1, followed by three consecutive ResNet3D blocks composed of two consecutive GroupNorm, SiLU non-linearity, and Conv3D with 3×3× 3 kernel, stride of 1 and padding of 1 and the resulting tensor is then added to the input (residual operation). After the first ResNet3D block, there is a Downsample3D block consisting of a Conv3D with kernel size of 3×3×3, stride of 2 and padding of 0. The three ResNet3D blocks in this stage gradually increase the number of filters (64, 128, and 256).

<u>***Stage 2*:**</u> Composed of a ResNet3D block followed by a CrossResolutionAttention3D - in which the feature map attends to a downsampled version of itself - followed by another ResNet3D block. The proposed CrossResolutionAttention3D creates attention maps between the raw input and its respective downsampled version by factor of 2 using MaxPool3d. Inspired by prior work (Adame-Gonzalez et al., 2023; Henschel et al., 2022) in which feature maps were downsampled using MaxPooling operations showing efficient and robust results, we propose this efficient modified self-attention mechanism that enables global patch attention operation using 8x less memory (i.e. theoretically 8x faster). The two ResNet3D blocks in this stage use the same number of filters (256).

<u>***Stage 3*:**</u> a GroupNorm followed by SiLU non-linearity and a Conv3D operation with 3×3×3 kernel, stride of 1 and padding of 1.

### 2.3.2. Decoder

**The generator decoder** *G*_*D*_ is similar in architecture to the encoder in reverse order (Figure 1-B):

***<u>Stage 3</u>:*** an initial Conv3D layer with stride of 1 and padding of 1 with a kernel size of 3×3×3 and 256 channels.

***<u>Stage 2</u>:*** two ResNet3D blocks with a CrossResolutionAttention3D block between them, with ResNet3D blocks using 256 channels each.

***<u>Stage 1</u>:*** Three ResNet3D blocks of decreasing channel count, with 256, 128, and 64 channels for the first, second, and third ResNet3D blocks, respectively. Between the second and third ResNet3D blocks, we apply an Upsample3D block that upsamples the 3D feature maps using nearest neighbor interpolation with an upsampling factor of 2.0 for all three axes to bring the dimensionality of the feature maps to match the output dimensions, followed by a Conv3D of 128 channels, kernel size of 3×3×3, padding and stride of 1. Finally, to generate the output image, we utilize a Conv3D unit with 1 output channel, kernel size of 3×3× 3, padding and stride of 1.

#### 2.3.3. Discriminator

**The discriminator** *D* is a patch-based discriminator (Demir & Unal, 2018; Rombach et al., 2022) consisting of 4 stages:

***<u>Stage 1</u>:*** initial Conv3D with 64 channels, kernel size of 4×4×4, stride of 2 and padding of 1 followed by a LeakyReLU non-linearity.

***<u>Stage 2</u>:*** a stack of 3 sequentially placed Conv3d with kernel size of 4×4×4, stride of 2 and padding of 1, followed by a BatchNorm3D layer and a LeakyReLU layer. In this stage, the first, second, and third Conv3d and BatchNorm3D operations have 128, 256, and 512 channels, respectively.

***<u>Stage 3</u>:*** A Conv3d with 512 filters, kernel size of 4×4×4, stride of 1 and padding of 1 followed by a BatchNorm3D and a LeakyReLU non-linearity.

***<u>Stage 4</u>:*** A single Conv3d layer with 512 input channels and 1 output channel using a kernel size of 4, with stride and padding of 1.

### 2.4. Training procedure

### 2.4.1. Generation of LR-HR image pair

Given a reference ground-truth 3D volume by the 5-dimensional tensor *x*∈*R*^*BS*×*C*×*H*×*W*×*D*^, where BS indicates the batch size, C indicates the number of channels (1 in our case), and H, W, and D the indicate dimensions of our 3D patch respectively (160, 160, 32), we define a set of random degradations by first defining our downsampling factors *df* = [*df*_*H*_, *df*_*W*_, *df*_*D*_], each of the three sampled from [1, 2, 3] with probabilities of [0.1, 0.45, 0.45] to ensure that downsampling factor of 1, i.e., no downsampling at all, is included in the training but with a small frequency compared to the other two downsampling factors. *df*_*H*_, *df*_*W*_, and *df*_*D*_ will be sampled independently during runtime. Then, an adaptive gaussian blurring function φ_*blur*_ will be applied by first defining the σ values for each axis in our image, i.e., defining σ = [σ_*H*_, σ_*W*_, σ_*D*_ ] with 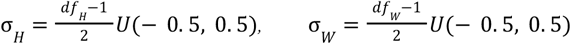, and 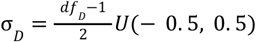, where *U*(− 0. 5, 0. 5) represents a uniform distribution and was added as a blurring augmentation. We then define the kernel size *k* for each axis as *k* = [*k*_*H*_, *k*_*W*_, *k*_*D*_ ], *k*_H_ = (2*ceil* (2σ_*H*_ + 1)), *k*_*W*_ = (2*ceil* (2σ_*W*_ + 1)), *k*_*D*_ = (2*ceil* (2σ_*D*_ + 1)) and apply the blurring independently to each axis using Conv3D from PyTorch. The mentioned blurring will be applied randomly to 50% of the training batches. Our low-resolution degraded image 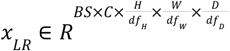 will be generated with *x*_*LR*_ = φ_↓,*df*_ φ_*lur,df*_ (*x*)_↓,*df*_ *w*here φ is a trilinear downsampling interpolation with downsampling factors 1/*df*. Finally, since our network design requires the input image to be in the same space (resolution) as the target image, we generate the final input to our network *x*_*LR*_*fake*_ ∈ *R*^*BS*×*C*×*H*×*W*×*D*^ with 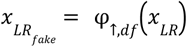 where φ_↑,*df*_ is a trilinear upsampling interpolation with factors *df*. The task of the network *G*_θ_ is to learn the mapping between *x*_*LR*_*fake*_ and *x*_*HR*_*fake*_ which is an approximation of *x*.

#### 2.4.2. Loss functions

*D* is optimized by minimizing *L* defined as

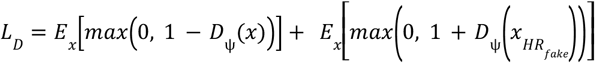 with *D* being the discriminator network parametrized by Ψ. The generator *G* is optimized by minimizing the loss function 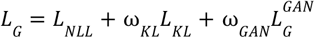, where *L*_*NLL*_ is defined as 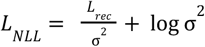. σ^2^ is a learnable parameter and *L*_*rec*_ is defined as *L*_*rec*_ = 2. 0*L*_*LPIPS*_ + 0. 2*L*_*MAE*_. Both *L*_*LPIPS*_ and *L*_*MAE*_ are defined as 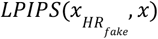 and 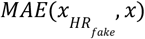. *L*_*KL*_ is the KL divergence between *z* defined as 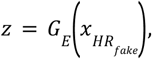 and a normal distribution *N*(0, *I*) defined as 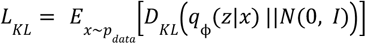, where q is the probability density mass of *G*_*E*_ parametrized by ϕ. 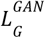 is defined as 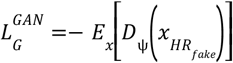, ω_*KL*_ is set to 1e-8, and ω_*GAN*_ is an adaptive function set to (Esser et al., 2021), 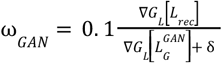 with ∇*G*_*L*_ [*L*_*rec*_ ] denotes the L2 norm of the gradients of *L*_*rec*_ with respect to the last layer of 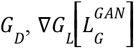 the L2 norm of the gradients of 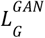 with respect to the last layer of *G*_*D*_, and δ is a stability constraint set to 10e-6.

### 2.4.3. Optimization

We trained RAVEN for 30 epochs, reaching convergence at epoch 22 (step ∼250k) using a batch size of 1, accumulating gradients for 2 iterations for G and 2 for D. ω_*KL*_ was set to 1e-8, and G was trained decoupled from D (G warmup) during one epoch. The learning rate was set to constant 9e-4 for the generator and 1.8e-3 for D, updating the discriminator 4 times per 1 update of the generator. Mixed-precision (bf16-mixed) was the datatype used for data efficiency, and the optimizer was set to AdamW with β1=0.5 and β2=0.9.

#### 2.4.4. Generating LR-HR image pairs

As stated above, few datasets contain images of the same participant/specimen acquired using the same scanner, same acquisition parameters, with the only difference being using two different resolutions (i.e., voxel sizes; either in-plane or slice-thickness). As such, it is necessary to generate LR surrogates that can closely resemble those acquired natively at LR. To achieve this, one could either apply k-space cropping (Forigua et al., 2022; Wu et al., 2023) (effectively removing the parts of the Fourier space corresponding to high-frequencies) or downsampling in the image domain using, e.g., interpolation after having blurred the image to prevent aliasing artifacts as described above (Zhang et al., 2021).

While the majority of proposed SISR methods report test results using the same degradation functions as those used for training, we tested the robustness of the trained network with the proposed degradation function φ_↓,*df*_ φ_*blur,df*_ to different degradation regimes such as interpolation after different intensities of gaussian blurring and k-space cropping. We opted to do so since no consensus exists on which degradation regime is best (i.e., most realistic) to generate low-resolution surrogates of an MRI for SISR tasks. Therefore, we assessed three most widely used test degradation schemes, two using varying parameters of φ_↓,*df*_ φ_*blur,df*_, and one using k-space cropping (Figure 3):

i. *<u>Degradation A:</u>* using φ_↓,*df*_ φ_*blur,df*_ defined for isotropic *df* values of [2, 2, 2] and [3, 3, 3]. σ = [σ_*H*_, σ_*W*_, σ_*D*_ ] are computed as 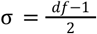.
ii. *<u>Degradation B :</u>* Same as *Degradation A*, but all σ were multiplied by a factor of 1.5, i.e., the blurring effect is more pronounced for this type of degradation.
iii. *<u>Degradation C:</u>* Given an input *x*∈*R*^*H*×*W*×*D*^ and a downsampling factor *df* = [*df*^*H*^, *df*^*W*^, *df*^*D*^], we apply the discrete Fourier transform *F* as ϖ = *F*(*x*) and perform the PyTorch *fftshift* (to bring the low-frequency components to the center-most locations of ϖ, obtaining ϖ_*shift*_). We then crop a volume of 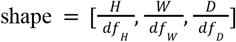 from the center-most elements of ϖ_*shift*_, obtaining ϖ_*LR*−*shift*_ and apply the inverse shifting operation, i.e., *ifftshift*. Finally, we generate the final downsampled image with the inverse Fourier transform as:

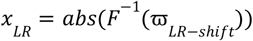

which is the magnitude of the complex voxel-wise volume returned by *F* ^−1^.

**Figure 3.**
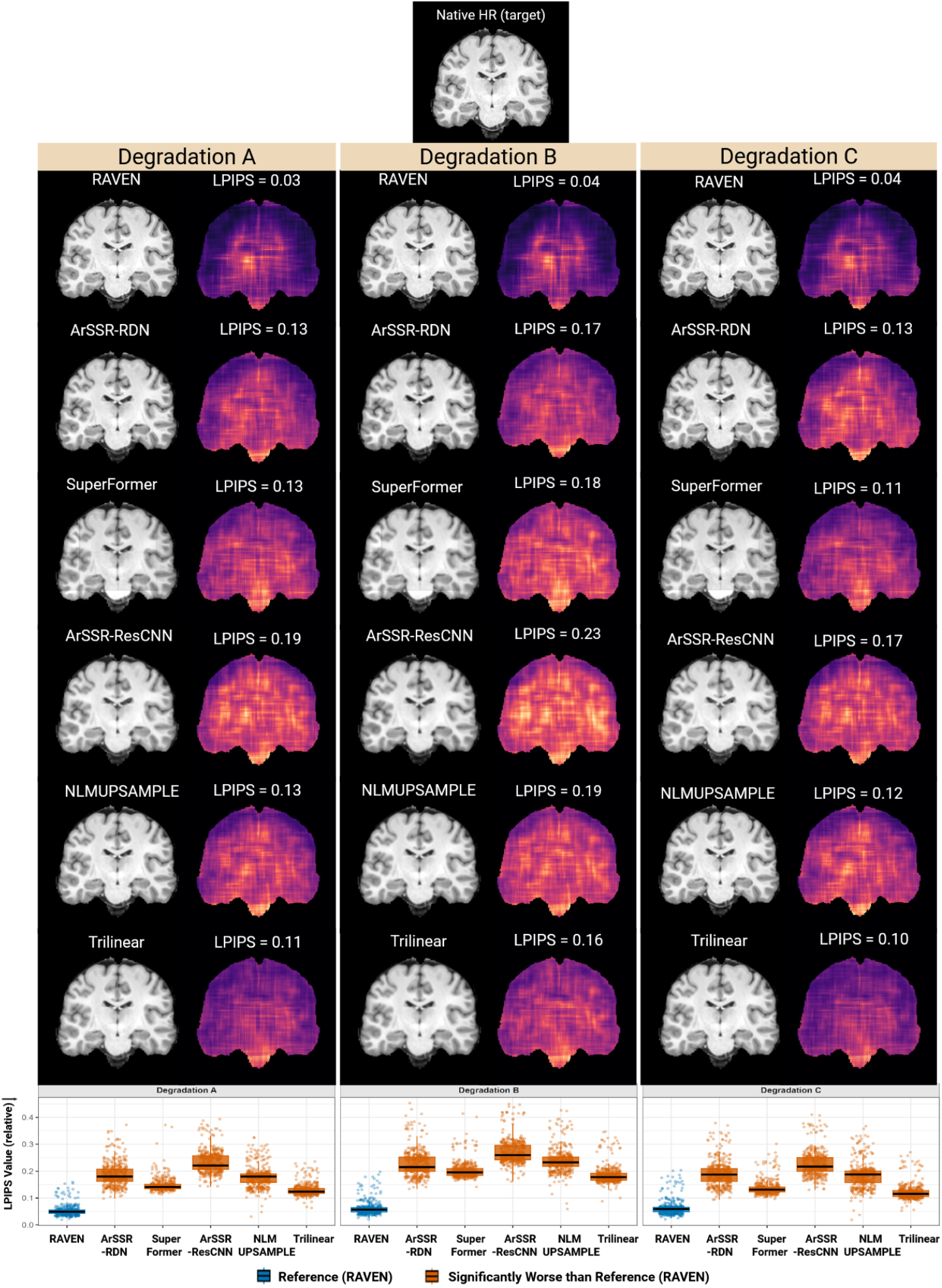
**Top:** Examples of coronal slices from a T1w image (the HCP dataset) downsampled by a factor of 2x isotropic using degradation schemes A, B, and C and then upsampled with different methods (left) as well as performance as measured by LPIPS (lower is better). **Bottom:** Boxplots reflecting the distribution of LPIPS values obtained for each method across the test dataset. Boxplot colors indicate significance of paired t-test vs. RAVEN after FDR correction. Each experiment was performed at both 2x and 3x isotropic upsampling factors.

**Figure 4.**
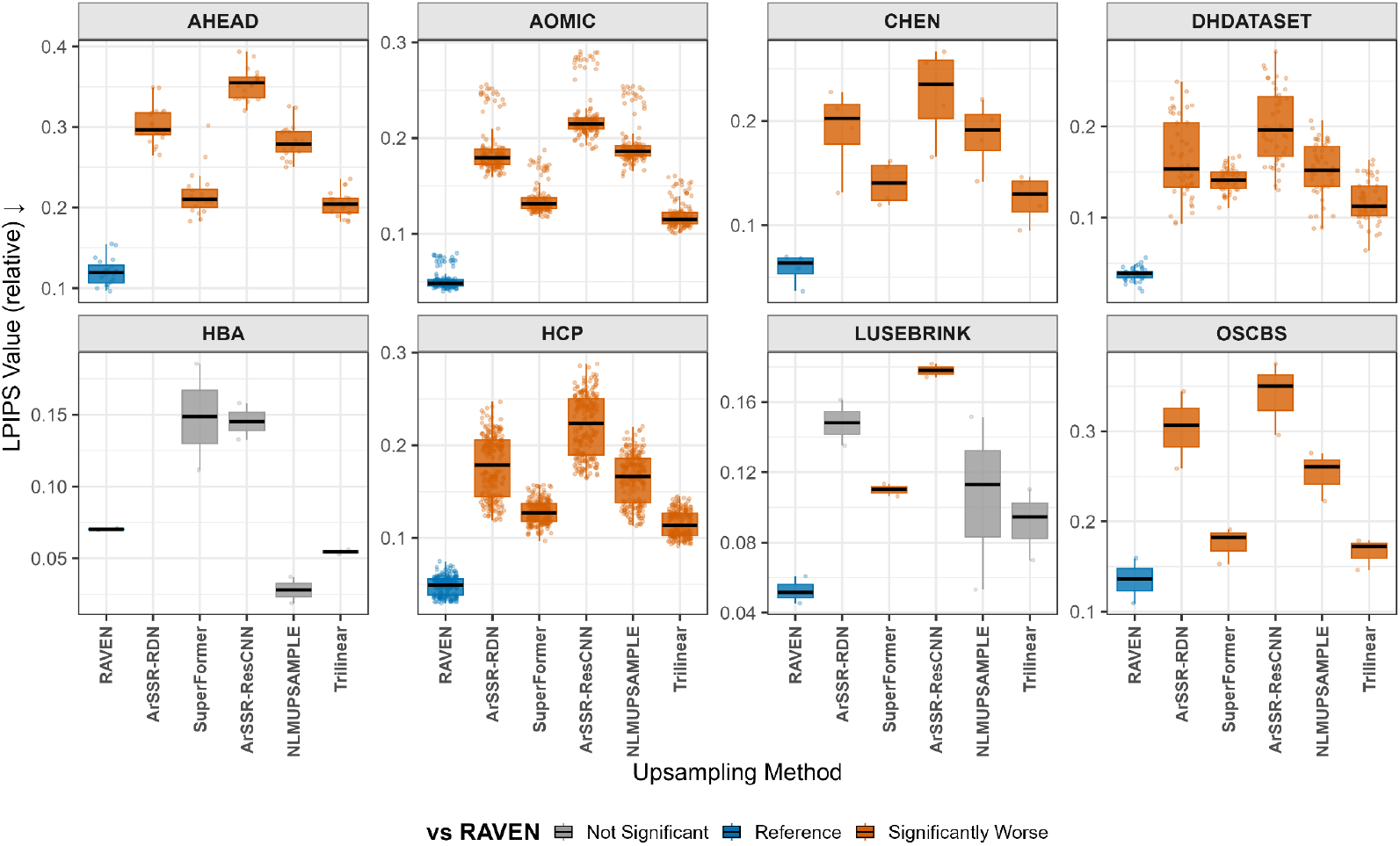
Performance per dataset as measured by LPIPS (lower is better). Boxplot colors indicate significance of paired t-test comparing other methods to RAVEN as the reference method following FDR correction. Each experiment was made at both 2x and 3x isotropic upsampling factors. Note that the lack of statistical significance in the differences for HBA and LUSEBRINK datasets is driven by their small sample sizes, rendering the tests and results unreliable.

## 3. Performance evaluation

The performance of RAVEN was compared against 4 state-of-the-art SISR methods, namely ArSSR-RDN (Wu et al., 2023), ArSSR-ResCNN (Wu et al., 2023), SuperFormer (Forigua et al., 2022), and NLMUPSAMPLE (Manjón et al., 2010), as well as trilinear interpolation. ArSSR-RDN and ArSSR-ResCNN were trained using convolutional neural networks, while SuperFormer is an implementation of Vision Transformers (ViT). NLMUPSAMPLE is a non-local patch image similarity method which iteratively optimizes the upsampled image until a convergence condition is met.

Mean Absolute Error (MAE), Peak Signal to Noise Ratio (PSNR), Structural Similarity Index Measure (SSIM), and Learned Perceptual Image Patch Similarity (LPIPS) (please refer to the appendix for details) were computed on the foreground mask by merging all the intracranial labels produced by SynthSeg (Billot et al., 2023), after resampling the results to the native image space using nearest neighbor interpolation. In the case of LPIPS and SSIM, we computed the slice-wise metrics averaged across the three anatomical planes.

Furthermore, to assess the impact of the methods on downstream derived measures, we generated SynthSeg masks from the upsampled MRIs and computed their similarity with the masks generated by the non-degraded native resolution HR image. We merged the bilateral Cerebral Gray Matter (GM), Cerebral White Matter (WM), Hippocampus, Ventricles, Deep GM, Cerebellum GMr, Cerebellum WM, and Brainstem to yield 8 ROIs for which we computed Dice (Dice, 1945) and IOU (Jaccard, 1901) similarity metrics. As all segmentations generated by SynthSeg have voxel sizes of 1mm isotropic, the resulting labels were resampled using nearest neighbor interpolator to match the space of the source MRIs prior to computing the segmentation agreement measures.

Paired t-tests were used to compare performance of methods for each metric, followed by false discovery rate (FDR) correction (Benjamini & Hochberg, 1995). Spearman’s and Pearson’s correlations were used to examine the associations between reconstruction performance metrics and segmentation performance.

## 4. Results

### 4.1. Across degradation schemes

RAVEN consistently outperformed all the tested methods on all four metrics, achieving lower LPIPS (See Figure 3) and MAE and higher PSNR and SSIM, for degradations A–C (see detailed stratified results in Table S2 and Figures S1 to S4). These results indicate that, irrespective of the degradation model, RAVEN yielded the most accurate and perceptually faithful reconstructions. Interestingly, Trilinear was consistently the second-best method in terms of LPIPS for upsampling factors of 2x, followed by NLMUPSAMPLE. For the rest of the metrics, NLMUPSAMPLE was the second-best method followed by Trilinear across all factors.

### 4.2. Across datasets

RAVEN was the top performer in the majority of cases. For LPIPS, RAVEN led on 7 out of 8 datasets (see detailed stratified results in Table S2 and Figures S1–4), tying with NLMUPSAMPLE on HBA. ArSSR-RDN was consistently second-best across degradations and datasets for upsampling factors of x3, while for factor of x2 the second-best method was Trilinear. For MAE, it led on 6 out of 8, tying with NLMUPSAMPLE, SuperFormer, and Trilinear on OSCBS and being outperformed by ArSSR-ResCNN, NLMUPSAMPLE, and Trilinear for HBA. For this metric, NLMUPSAMPLE was consistently the second-best method; for PSNR, RAVEN led on 7 out of 8, with HBA favoring Trilinear, NLMUPSAMPLE, and ArSSR-ResCNN; and for SSIM, RAVEN outperformed all other methods across datasets except for HBA, in which NLMUPSAMPLE and Trilinear outperformed RAVEN at x2 upsampling factors.

### 4.3. By target voxel size

Stratifying by voxel size (Table S3; Figures S5-8) further supported RAVEN’s robustness. RAVEN led LPIPS across all bins (≤0.60 mm, 0.60–0.90 mm, >0.90 mm) (see Figure 5), and achieved the best MAE in two out of three bins while matching NLMUPSAMPLE and Trilinear at ≤0.60 mm. For PSNR, RAVEN dominated at 0.60–0.90 mm and >0.90 mm, whereas NLMUPSAMPLE led at ≤0.60 mm. For SSIM, RAVEN led across all voxel bins (see detailed stratified results on Figures S5–8).

**Figure 5.**
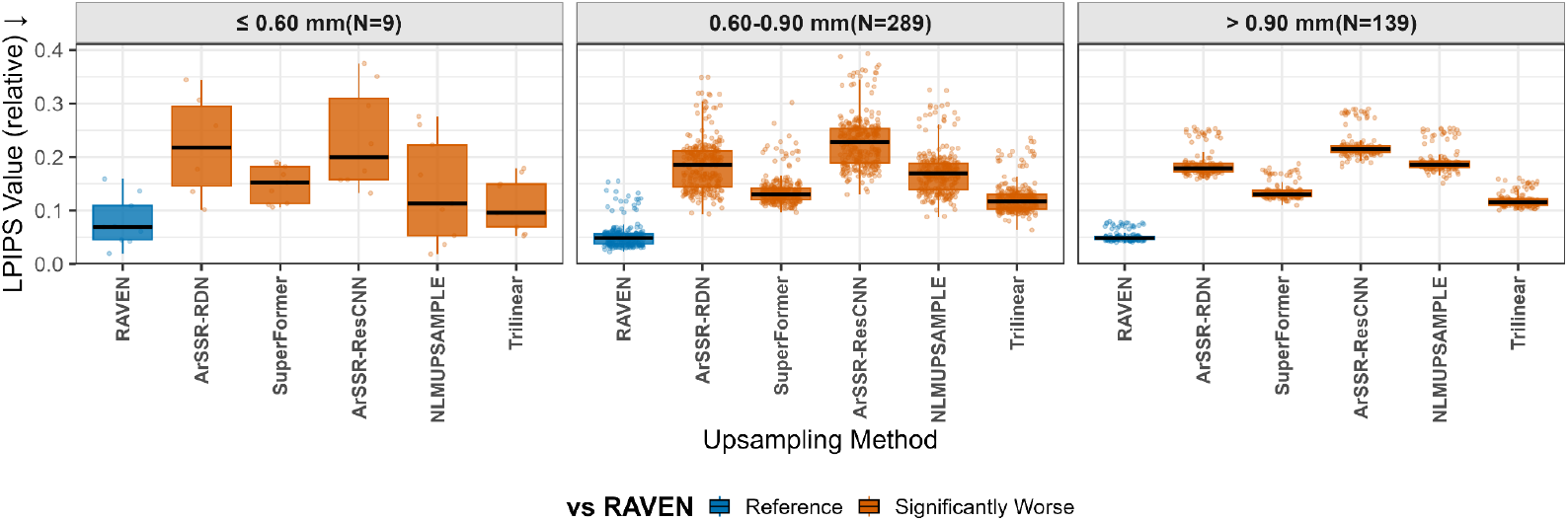
Performance per dataset as measured by LPIPS (lower is better). Boxplot colors indicate significance of paired t-tests comparing other methods to RAVEN as the reference method following FDR correction. Each experiment was made at both 2x and 3x isotropic upsampling factors.

### 4.4. Downstream segmentation of upsampled MRIs

RAVEN yielded the best results across metrics, upsampling factors, datasets, degradation methods, and ROIs with 0.906 and 0.870 DICE and IOU, followed by Trilinear interpolation with 0.880 and 0.835, and SuperFormer with 0.856 and 0.801 DICE and IOU respectively. ArSSR (both resCNN and RDN versions) yielded the lowest segmentation agreement metrics with DICE of 0.815 and IOU of 0.73 for both ArSSR-ResCNN and ArSSR-RDN. For all 8 regions of interest, whole-brain LPIPS was the image similarity metric that better predicted segmentation agreement with the reference in terms of both Spearman’s and Pearson’s correlations (See Figure 6 as well as Figure S10).

**Figure 6.**
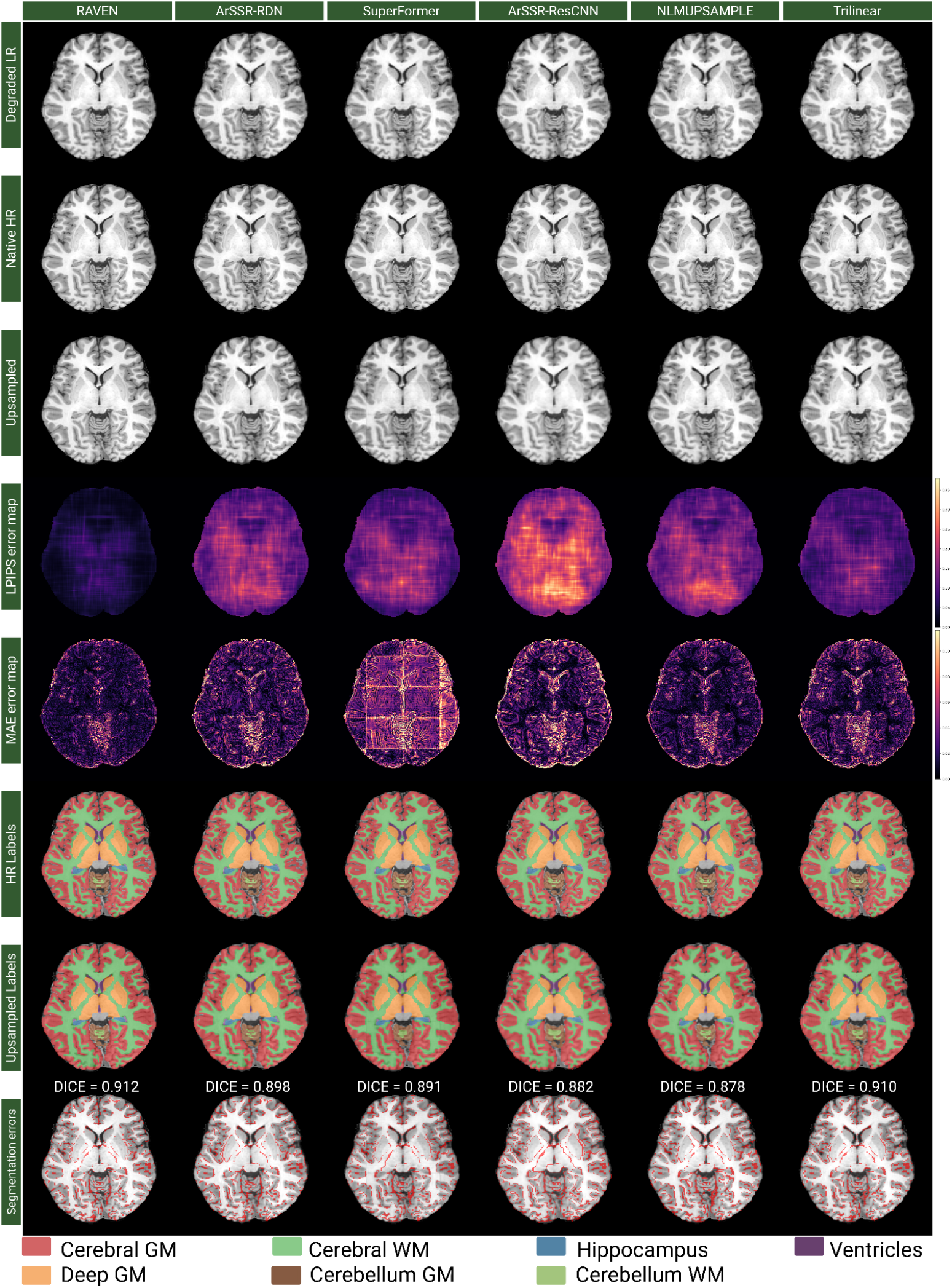
Example of an image from the HCP dataset degraded using Degradation scheme C and downsampling factor of 2 on all tested super-resolution methods. From top to bottom rows: degraded image (Native LR), non-degraded native image (Native HR), upsampled image (Native LR upsamped to Native HR space), LPIPS error map produced between Native HR and Upsampled rows, MAE error map (difference between Native HR and Upsampled rows), HR labels (SynthSeg masks merged into 8 ROIs), Upsampled LR Labels (SynthSeg masks from the Upsampled row), Difference between HR and Upsampled LR masks.

### 4.5. From true LR to true HR data

To test the proposed method with ground-truth LR-HR image pairs, we acquired images of stabilized post-mortem head and hemisphere specimens during the same scanning session using the same parameters in LR and HR (e.g. T2w scans acquired at both 0.64mm and 1.28mm isotropic). We upsampled the true LR image using both NLMUPSAMPLE and RAVEN from 1.28mm isotropic to 0.64mm isotropic. We then computed the MAE, PSNR, SSIM, and LPIPS on the foreground after merging the intracranial labels produced by SynthSeg (Billot et al., 2023). RAVEN outperformed NLMUPSAMPLE across all metrics, with MAE, PSNR, SSIM, and LPIPS values of 0.048, 30.68dB, 0.741, and 0.176 respectively compared to 0.056, 29.28dB, 0.736, and 0.253 obtained for NLMUPSAMPLE.

Finally, to extract MRI-derived metrics after applying super-resolution, we applied SynthSeg (Figure 7) and generated 8 regions of interest (ROI): Cerebral and cerebellar Gray and White matter, hippocampus, ventricles, deep Gray matter, and brainstem. We observed consistent results across the performed tests: RAVEN obtained the best performance across ROIs with mean DICE and IOU of 0.91 (±0.03) and 0.83 (±0.05) respectively, followed by trilinear interpolation with 0.90 (±0.031) and 0.83 (±0.052) and then SuperFormer with 0.89 (±0.032) and 0.81 (±0.051) DICE and IOU respectively.

**Figure 7.**
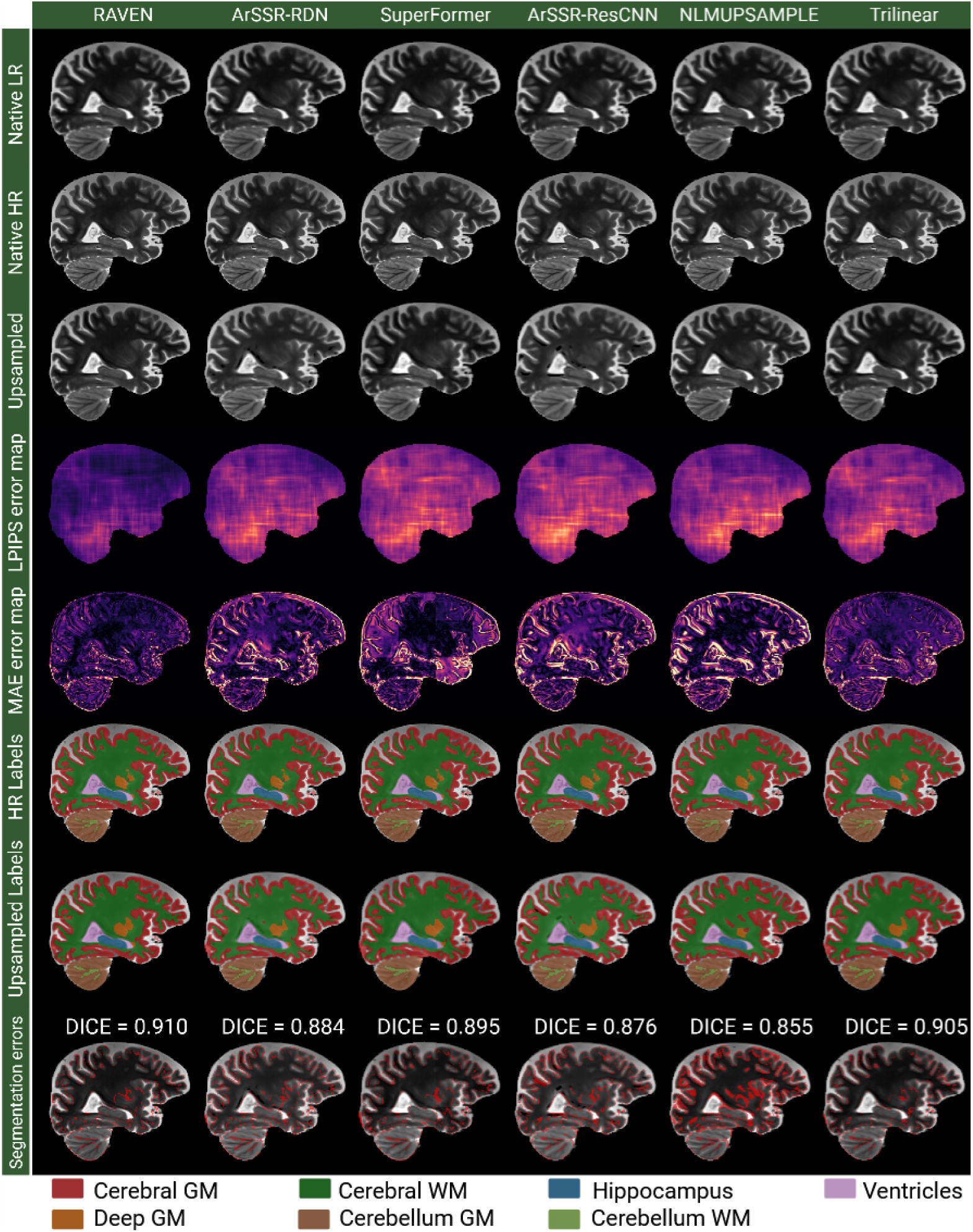
Performance of all tested methods with gold-standard high-resolution ground truth on unseen full-head post-mortem T2w MRI. RAVEN achieves highest similarity which becomes

## 5. Discussion

In this work, we presented RAVEN, an open source, resolution and contrast agnostic, SISR method. We demonstrated that RAVEN obtains competitive results when compared against other state-of-the-art methods across degradation modes, resolutions, and upsampling factors. RAVEN also outperformed other methods with respect to downstream task performance: the segmentations generated by SynthSeg on the upsampled volumes were more similar to the reference ground truth than those obtained with interpolation. RAVEN obtained higher overall segmentation performance across metrics, downsampling factors, degradation modes, and datasets, indicating higher fidelity in tissue reconstruction in high-resolution domains.

Despite being a 3D model, RAVEN’s patch-based inference design allows it to run on domestic-grade GPUs with less than 24GB. Additionally, to limit the peak VRAM usage through the patch-based inference, the inclusion of the CrossResolutionAttention3D module allows for a pseudo-self-attention operation for global context inclusion using 8x less memory, making it a suitable option for its usage in contexts where high-performance clusters are out of reach.

Degradation C (k-space cropping) follows MRI-physics more closely than gaussian blurring followed by downsampling and should be used as a degradation function for generating LR-HR image pairs. While the first statement is true, there is evidence in the SISR literature (Rombach et al., 2022; Zhang et al., 2021) that supports the usage of a diverse set of degradations with multiple blurring and downsampling operations in favor of better generalizability. We posit that simply cropping the k-space as it is done in recent literature (Wu et al., 2023; K. Zhao et al., 2025) does not provide sufficient input variability. These observations are supported by the results reported in this work, in which our simple but varied set of degradations applied during training enabled a well-behaved network when dealing with unseen degradations as shown by the performance metrics on Degradation C.

While ArSSR-RDN (Wu et al., 2023), ArSSR-ResCNN (Wu et al., 2023), and SuperFormer (Forigua et al., 2022) are recent state-of-the-art methods, only ArSSR-RDN consistently improved reconstruction over trilinear interpolation among the tested deep-learning algorithms. This improvement, however, was observed only at 3×3×3 upsampling factors. This is likely since the method was trained using upsampling factors in the range of [2-4] and performed better at 3×3×3 (i.e. mean of the selected range) compared to 2×2×2 in the results reported in the original article as well. We hypothesize that optimizing for voxel-space metrics and using a single dataset with only T1w images are detrimental to generalizability across other datasets and upsampling factors.

All metrics were carefully quality-controlled, and our procedure was adapted to produce robust measurements: only voxels corresponding to brain tissue were considered; robust outlier cropping was applied uniformly to all images (clamping intensities to the 0.5th and 99.5th percentiles); and LPIPS and SSIM were computed as the average across the three anatomical planes. This procedure prevents the metrics from being severely affected by outliers.

Another relevant contribution of this work is demonstrating that LPIPS is a superior image-quality metric that better detects true structural differences than voxel-space, intensity-based metrics such as PSNR and MAE. LPIPS showed the highest correlation with DICE and IOU, followed by SSIM, in our downstream segmentation tests with SynthSeg (Figure S.10). Although RAVEN did not outperform all methods across all scenarios for MAE or PSNR, it achieved the best LPIPS and the highest IOU and DICE. These findings support the view that deep-feature similarity metrics and loss functions better preserve true anatomy, and suggest that a future 3D-LPIPS distance for medical-image applications could further improve robustness.

Our loss function was designed with fine-detail reconstruction in mind (small structures, high-frequency components); thus, LPIPS was the dominant component of the reconstruction loss. An L1 term was added to prevent shifts in the overall intensity distribution. Because the network uses 3D patches while LPIPS is inherently 2D, we adapted the LPIPS component to compute a loss for each anatomical plane and average the three per-plane values, regularizing the network and preventing slicing artifacts that could lead to a “jagged” appearance on perpendicular planes. GAN training is known to present multiple challenges, especially with training dynamics. The best stability was achieved by training the generator for two epochs without a discriminator and then introducing the discriminator. Additionally, tuning the contribution of the adaptive weight was the second most crucial factor for a well-behaved, convergent network.

Note that all generated results for all networks were obtained using the same set of author-released pretrained weights and no test-specific retraining was performed. Thus, the generalization capability of a single set of weights per method was tested across all resolutions and MRI modalities described above. Furthermore, ArSSR-RDN (Wu et al., 2023), ArSSR-ResCNN (Wu et al., 2023), and SuperFormer (Forigua et al., 2022) were trained and validated based on the HCP dataset in the original articles, and as such, the results based on the HCP dataset reflect their within sample performance, and the other datasets can be considered out of sample. Importantly, even in this within sample comparisons, RAVEN outperformed all these models. We provided examples of the methods’ performances for 2 HCP images in Figures 3 and 6, to demonstrate RAVEN’s superior performance even in the dataset the other methods had specialized on. Additionally, Table S2 and Figs S1-S4 provide detailed information about the results obtained per method and further stratified by experiment (dataset, degradation mode, and upsampling factor).

This work is not without limitations. The most evident limitation is the lack of substantial numbers of true LR–HR image pairs acquired at native low and high resolutions, which would be ideal for network training. Using the proposed degradation functions during training was one strategy to boost RAVEN’s generalization performance. If one had access to a sufficiently large, varied set of native LR–HR MRI pairs, one could train a GAN that generates degradations to better simulate LR MRIs from HR counterparts and then train a SISR network using those synthetic outputs. Another limitation is model size: larger models typically perform better. Although our environment would allow training larger models, and despite our CrossResolutionAttention3D enabling a more efficient attention mechanism, we opted to design a model that can be trained locally on an NVIDIA RTX 4090 GPU—a relatively low-cost, non-research-grade GPU—to optimize resources, enable efficient deployment by most medical-image processing research groups worldwide, and promote access to state-of-the-art tools in low-income countries. We therefore acknowledge that a version of RAVEN with more filters per convolutional layer would likely achieve higher performance in less-constrained environments.

Additionally, we recognize that alternative state-of-the-art methods for MRI SISR exist. The denoising diffusion probabilistic model (DDPM) family has gained popularity as it has demonstrated the ability to outperform GANs and has been applied to brain MRI SISR (J. Wang et al., 2023; K. Zhao et al., 2025)). However, these models typically require substantial computational resources, often operate on 2D slices, may not directly increase in-plane resolution, and, by design, rely on iterative inference along an inverse diffusion Markov chain. An alternative that could enable faster inference is latent flow matching (Fischer et al., 2024). Exploring such iterative algorithms remains future work.

In this work, we proposed and validated RAVEN, a robust, generalizable, contrast-agnostic, and efficient deep-learning method for brain MRI SISR. RAVEN is open-source with open-access weights, and can be run in non-research grade GPUs. We extensively validated RAVEN on in-vivo and ex-vivo datasets acquired at a wide range of voxel sizes and modalities. Furthermore, we assessed the downstream quality of segmenting the upsampled MRIs, demonstrating that RAVEN achieves state-of-the-art performance, significantly improving over other tested super-resolution methods.

## 6. Acknowledgements

We gratefully acknowledge the open datasets that made this work possible: AHEAD (Alkemade et al., 2020); the Amsterdam Open MRI Collection — ID1000, PIOP1, and PIOP2 (Snoek et al., 2021); the paired 3T/7T T1/T2 dataset (X. Chen et al., 2023); HumanBrainAtlas (Schira et al., 2023); the Open Science CBS Neuroimaging Repository (Tardif et al., 2016); the ultrahigh-resolution 7 T dataset of (Lüsebrink et al., 2017, 2021); and post-mortem MRI samples provided by the Douglas-Bell Canada Brain Bank—we especially thank the donors and their families (Dadar et al., 2024). Data were provided [in part] by the Human Connectome Project, WU-Minn Consortium (Principal Investigators: David Van Essen and Kamil Ugurbil; 1U54MH091657) funded by the 16 NIH Institutes and Centers that support the NIH Blueprint for Neuroscience Research; and by the McDonnell Center for Systems Neuroscience at Washington University.

## 7. Ethics

Ethics approval for all datasets used was obtained at respective sites.

## 8. Funding

Dadar reports receiving research funding from the Healthy Brains for Healthy Lives (HBHL), Alzheimer Society Research Program (ASRP), Douglas Research Centre (DRC), and Natural Sciences and Engineering Research Council of Canada (NSERC), Canadian Institutes of Health Research (CIHR), Fonds de Recherche du Quebec—Santé (FRQS, DOI https://doi.org/10.69777/330750), and Brain Canada. Zeighami reports receiving research funding from the Healthy Brains for Healthy Lives, Fonds de recherche du Québec—Santé (FRQS, DOI https://doi.org/10.69777/320107) Chercheurs boursiers et chercheuses boursières en Intelligence artificielle, as well as Natural Sciences and Engineering Research (NSERC) discovery grant. Adame-Gonzalez reports receiving funding from Fonds de recherche du Québec—Santé (FRQS, DOI https://doi.org/10.69777/351384).

## 9. Author Contributions

W.A.-G.: study design, data preprocessing, coding, testing, analysis of results, figure generation, manuscript writing R.M.: Data curation, quality control. Y.Z.: Study design, conceptualization, writing, analysis of results, project supervision, funding. M.D.: Study design, conceptualization, writing, analysis of results, project supervision, funding.

## Supplementary Materials

### Data Description

#### Amsterdam Ultra-high field adult lifespan database (AHEAD)

AHEAD (Alkemade et al., 2020) was designed to characterize healthy adult lifespan variation at ultra-high field with submillimeter, quantitative 7T MRI optimized for subcortical anatomy.

#### Amsterdam Open MRI Collection (AOMIC)

Comprises three main datasets (Snoek et al., 2021). **ID1000** is part of the Amsterdam Open MRI Collection and was designed as a large normative 3T cohort to benchmark methods and study variability in brain–behavior relationships. **PIOP1** targets task-based individual-differences mapping in young adults to support robust, reproducible cognitive-neuroimaging pipelines. **PIOP2** complements PIOP1 with additional tasks and phenotypes to facilitate replication and modeling of brain–behavior links under harmonized 3 T acquisitions.

#### CHEN

This paired 3T/7T T1w/T2w dataset was created to enable development and evaluation of 3T to 7T image synthesis methods using matched contrasts within subjects (X. Chen et al., 2023).

#### DHDATASET

Developed to standardize post-mortem MRI acquisition in a large brain bank and to link MRI with histology for neurodegenerative disease research (Dadar et al., 2024).

#### Human Brain Atlas (HBA)

HBA was designed to provide ultra-detailed in-vivo MRI with expert segmentations for fine-scale neuroanatomy, and method validation at very high spatial resolution (Schira et al., 2023).

#### Human Connectome Project (HCP)

HCP established high-quality 3T MRIs in healthy young adults to map structural and functional connectivity and serve as a community benchmark for pipelines/analyses.

#### Open Science CBS Neuroimaging Repository (OSCBS)

OSCBS was created to disseminate ultra-high-field (7T) quantitative/anatomical datasets (e.g., MP2RAGE, multi-echo FLASH) for high-resolution method development and validation (Tardif et al., 2016).

#### LUSEBRINK

Proposed as an ultra-high-resolution MRI “human phantom”, this dataset was designed to test feasibility of fine-scale morphometry and algorithm testing with 7T in-vivo anatomy (Lüsebrink et al., 2017).

---

**Table S1.** Performance of tested methods per degradation type and upsampling factor (scale). Metrics reported as “mean ± (standard deviation)”.

| Degradation | Scale | Upsampling Method | LPIPS ↓ | MAE ↓ | PSNR ↑ | SSIM ↑ |
| --- | --- | --- | --- | --- | --- | --- |
| Degradation A | 2x2x2 | RAVEN | 0.052 ± (0.019) | 0.030 ± (0.015) | 35.860 ± (3.182) | 0.900 ± (0.040) |
| | | ArSSR-RDN | 0.187 ± (0.052) | 0.047 ± (0.029) | 31.121 ± (3.557) | 0.824 ± (0.061) |
| | | SuperFormer | 0.147 ± (0.025) | 0.058 ± (0.029) | 29.359 ± (3.638) | 0.826 ± (0.046) |
| | | ArSSR-ResCNN | 0.232 ± (0.040) | 0.054 ± (0.022) | 29.755 ± (3.397) | 0.782 ± (0.049) |
| | | NLMUPSAMPLE | 0.179 ± (0.039) | 0.036 ± (0.018) | 34.330 ± (3.188) | 0.873 ± (0.041) |
| | | Trilinear | 0.130 ± (0.021) | 0.039 ± (0.016) | 33.827 ± (3.098) | 0.851 ± (0.036) |
| | 3x3x3 | RAVEN | 0.149 ± (0.038) | 0.044 ± (0.022) | 32.120 ± (3.294) | 0.831 ± (0.047) |
| | | ArSSR-RDN | 0.282 ± (0.047) | 0.073 ± (0.024) | 27.181 ± (2.679) | 0.662 ± (0.055) |
| | | SuperFormer | 0.339 ± (0.037) | 0.080 ± (0.024) | 26.271 ± (2.779) | 0.662 ± (0.045) |
| | | ArSSR-ResCNN | 0.336 ± (0.043) | 0.080 ± (0.026) | 26.300 ± (2.851) | 0.620 ± (0.052) |
| | | NLMUPSAMPLE | 0.294 ± (0.044) | 0.055 ± (0.021) | 30.390 ± (2.890) | 0.750 ± (0.044) |
| | | Trilinear | 0.333 ± (0.040) | 0.068 ± (0.023) | 28.195 ± (2.643) | 0.680 ± (0.045) |
| Degradation B | 2x2x2 | RAVEN | 0.060 ± (0.023) | 0.030 ± (0.017) | 35.630 ± (3.397) | 0.908 ± (0.038) |
| | | ArSSR-RDN | 0.226 ± (0.045) | 0.053 ± (0.022) | 29.825 ± (3.402) | 0.806 ± (0.050) |
| | | SuperFormer | 0.202 ± (0.030) | 0.063 ± (0.026) | 28.578 ± (3.362) | 0.793 ± (0.043) |
| | | ArSSR-ResCNN | 0.271 ± (0.044) | 0.060 ± (0.024) | 28.816 ± (3.333) | 0.758 ± (0.049) |
| | | NLMUPSAMPLE | 0.240 ± (0.057) | 0.045 ± (0.034) | 32.273 ± (3.388) | 0.833 ± (0.056) |
| | | Trilinear | 0.185 ± (0.031) | 0.047 ± (0.020) | 31.707 ± (3.073) | 0.814 ± (0.040) |
| | 3x3x3 | RAVEN | 0.207 ± (0.043) | 0.055 ± (0.024) | 29.920 ± (3.079) | 0.771 ± (0.050) |
| | | ArSSR-RDN | 0.345 ± (0.044) | 0.079 ± (0.025) | 26.667 ± (2.590) | 0.623 ± (0.054) |
| | | SuperFormer | 0.425 ± (0.042) | 0.092 ± (0.026) | 24.919 ± (2.592) | 0.596 ± (0.046) |
| | | ArSSR-ResCNN | 0.409 ± (0.041) | 0.085 ± (0.026) | 25.933 ± (2.579) | 0.578 ± (0.053) |
| | | NLMUPSAMPLE | 0.388 ± (0.047) | 0.069 ± (0.023) | 27.999 ± (2.623) | 0.674 ± (0.046) |
| | | Trilinear | 0.423 ± (0.045) | 0.079 ± (0.023) | 26.557 ± (2.400) | 0.614 ± (0.047) |
| Degradation C | 2x2x2 | RAVEN | 0.062 ± (0.026) | 0.030 ± (0.015) | 35.482 ± (3.190) | 0.901 ± (0.038) |
| | | ArSSR-RDN | 0.189 ± (0.046) | 0.046 ± (0.019) | 31.392 ± (3.153) | 0.824 ± (0.049) |
| | | SuperFormer | 0.137 ± (0.027) | 0.056 ± (0.022) | 29.438 ± (3.223) | 0.829 ± (0.040) |
| | | ArSSR-ResCNN | 0.227 ± (0.044) | 0.052 ± (0.021) | 30.165 ± (3.280) | 0.791 ± (0.048) |
| | | NLMUPSAMPLE | 0.185 ± (0.045) | 0.034 ± (0.016) | 34.810 ± (3.187) | 0.877 ± (0.040) |
| | | Trilinear | 0.122 ± (0.028) | 0.040 ± (0.017) | 33.371 ± (3.042) | 0.845 ± (0.042) |
| | 3x3x3 | RAVEN | 0.177 ± (0.041) | 0.045 ± (0.019) | 32.003 ± (2.912) | 0.796 ± (0.048) |
| | | ArSSR-RDN | 0.273 ± (0.047) | 0.069 ± (0.022) | 27.794 ± (2.463) | 0.657 ± (0.053) |
| | | SuperFormer | 0.309 ± (0.039) | 0.073 ± (0.023) | 27.163 ± (2.670) | 0.690 ± (0.046) |
| | | ArSSR-ResCNN | 0.305 ± (0.044) | 0.073 ± (0.024) | 27.176 ± (2.719) | 0.637 ± (0.051) |
| | | NLMUPSAMPLE | 0.277 ± (0.048) | 0.049 ± (0.018) | 31.292 ± (2.773) | 0.765 ± (0.046) |
| | | Trilinear | 0.305 ± (0.042) | 0.061 ± (0.021) | 29.287 ± (2.697) | 0.704 ± (0.047) |

**Table S2.**
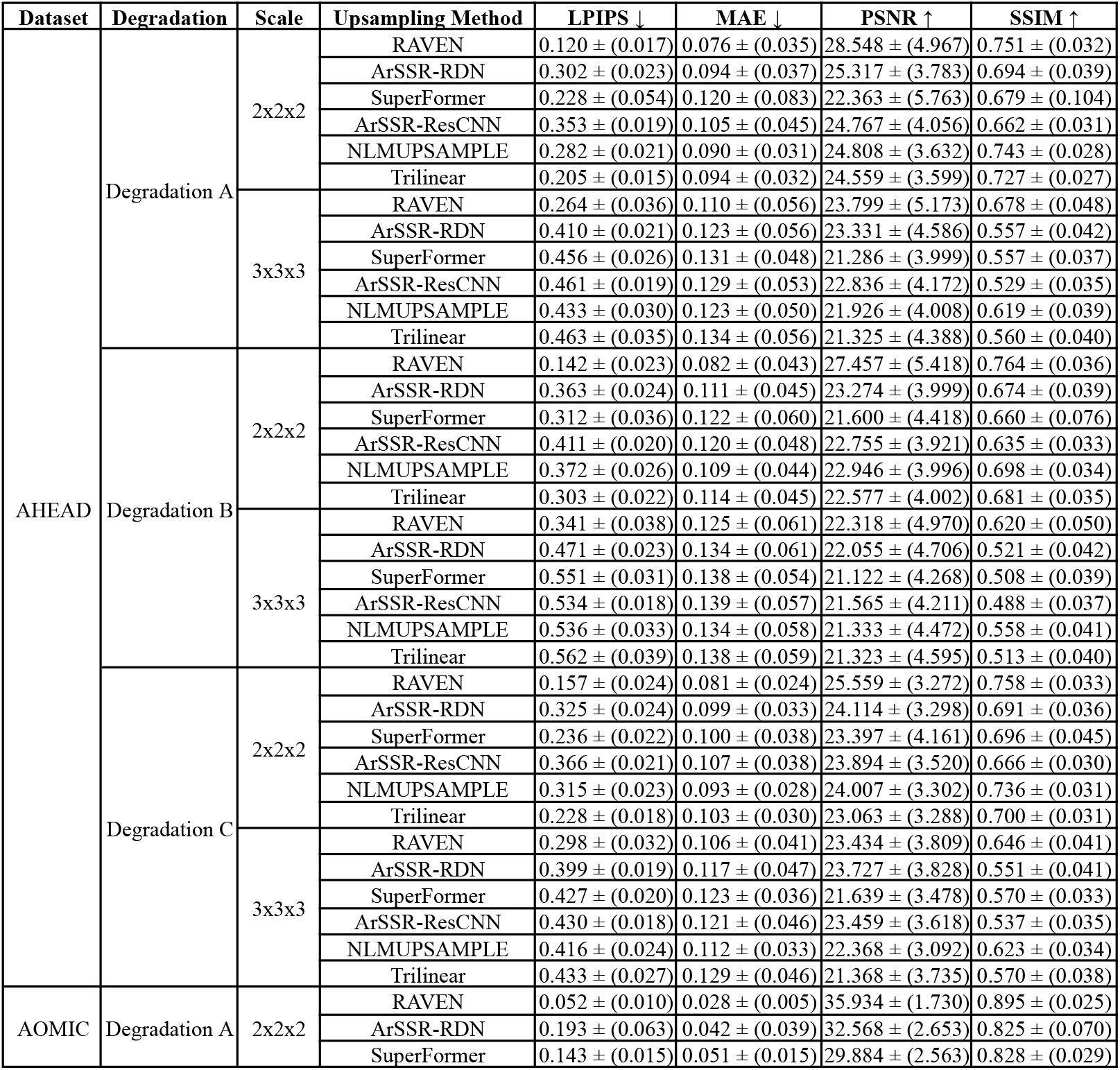
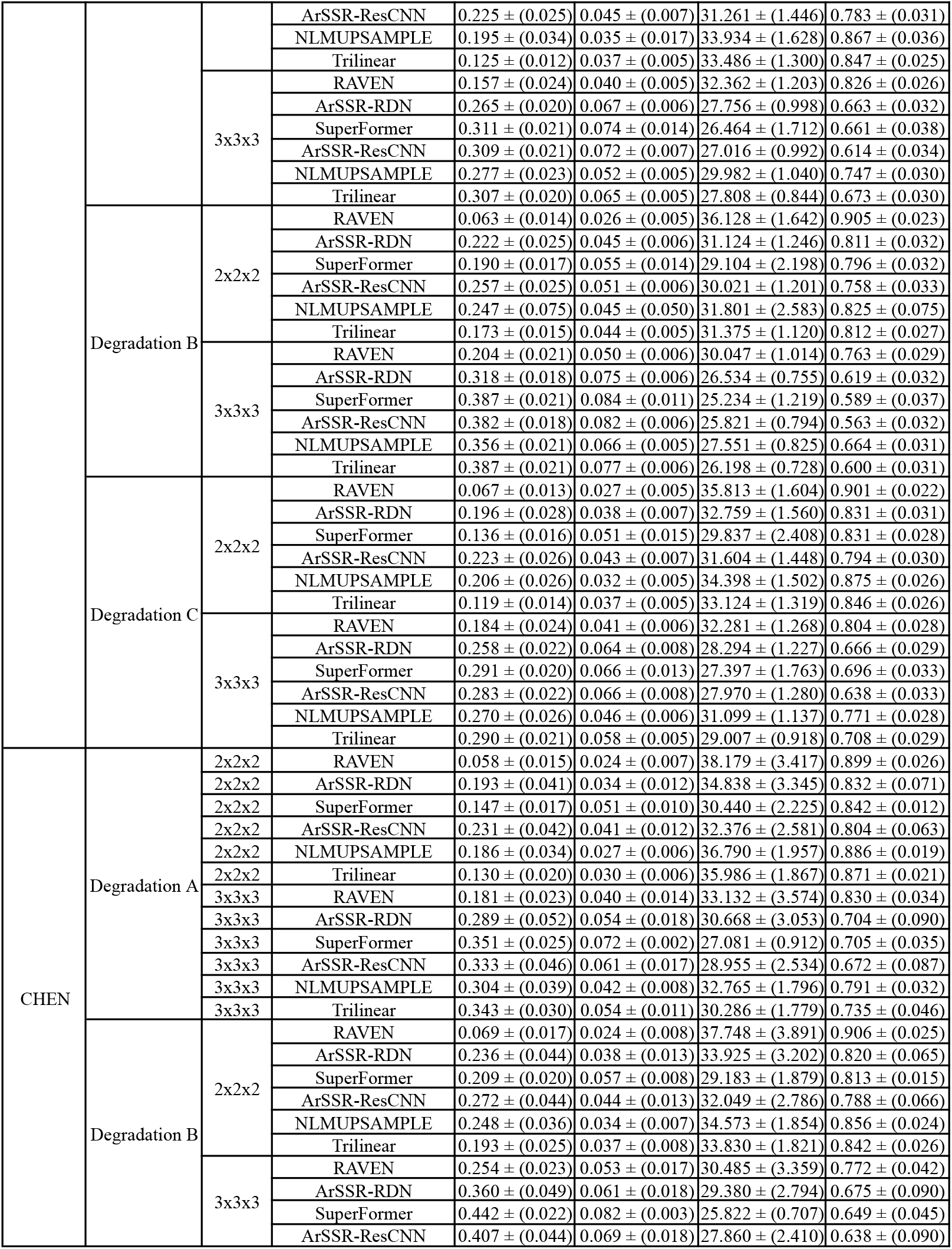
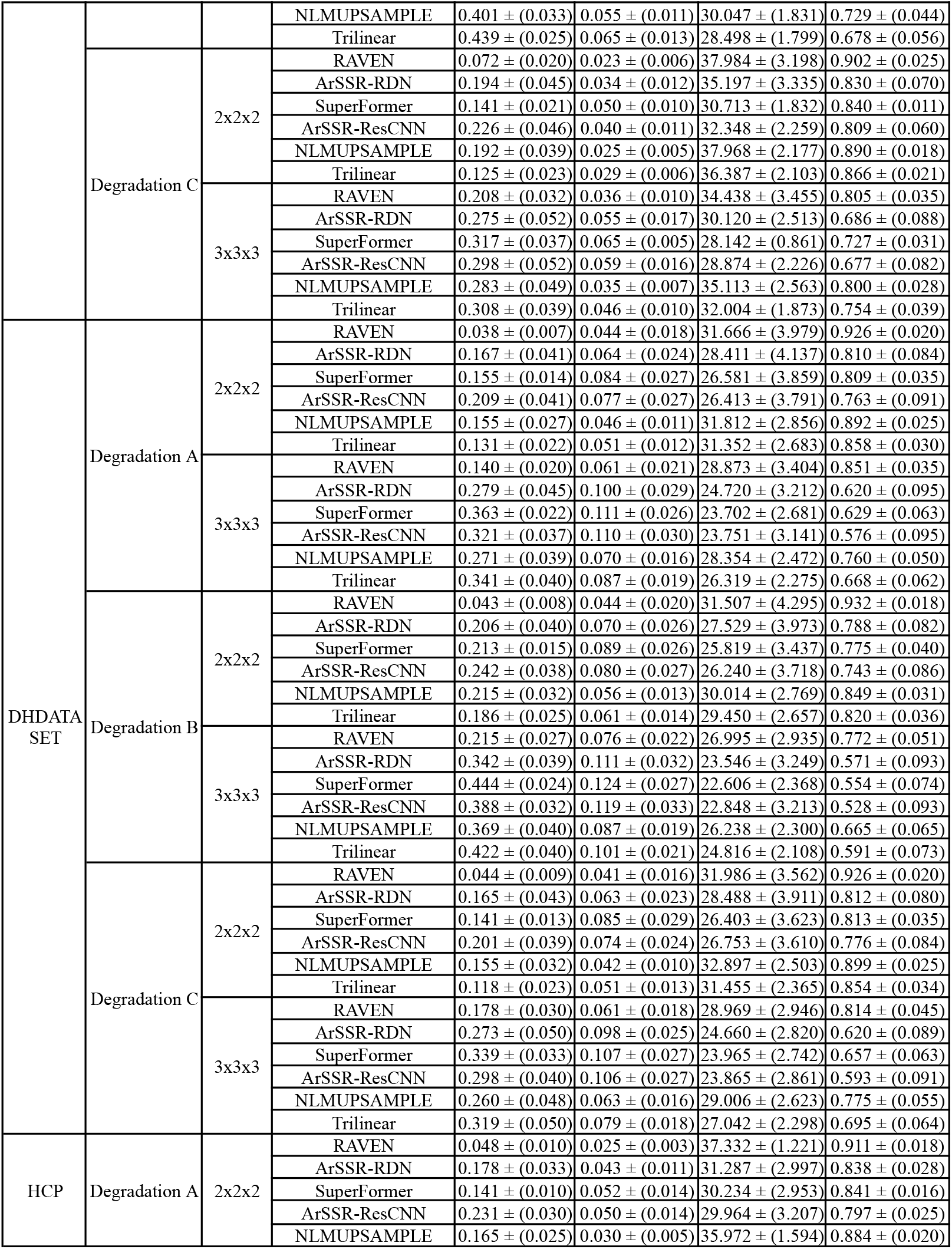
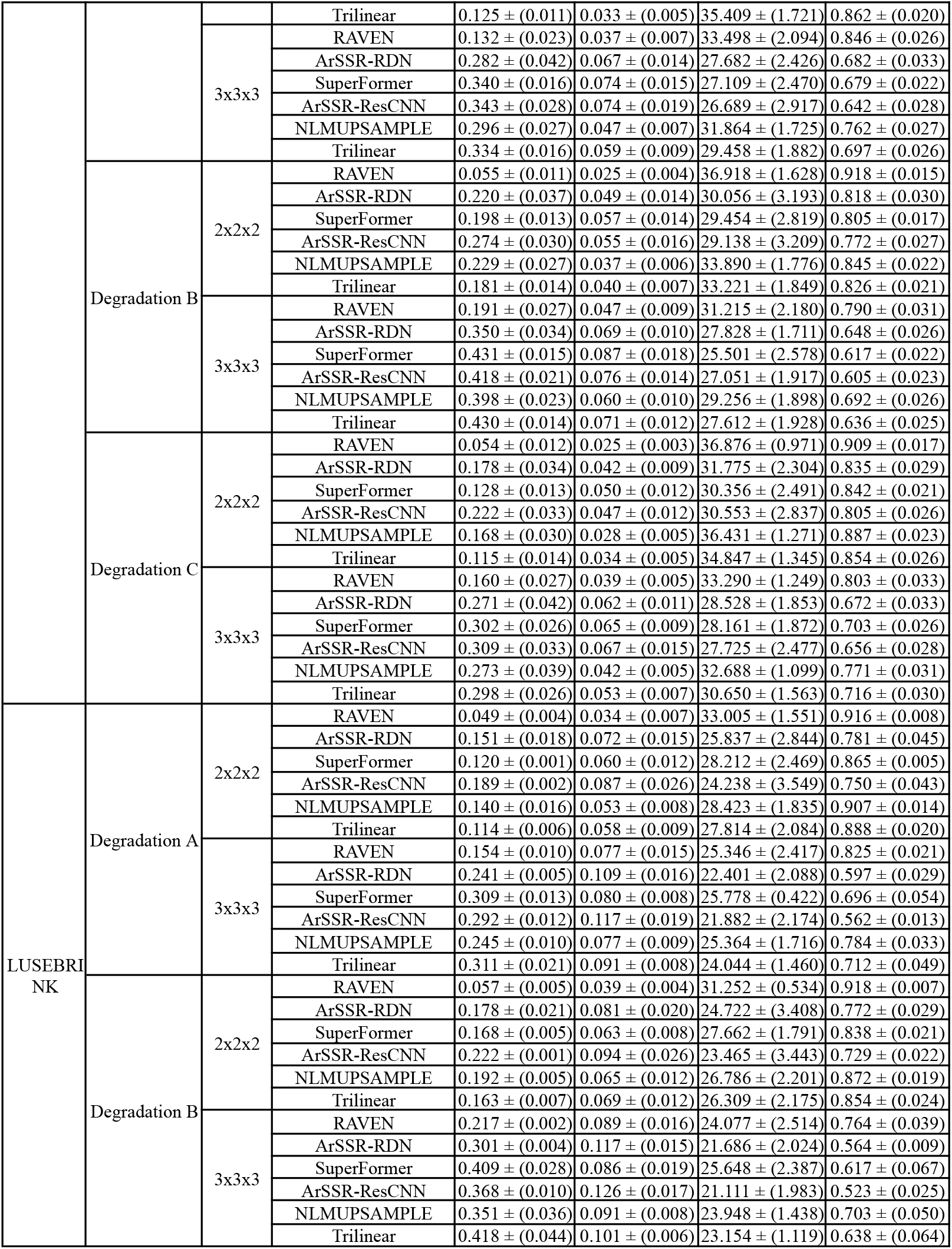
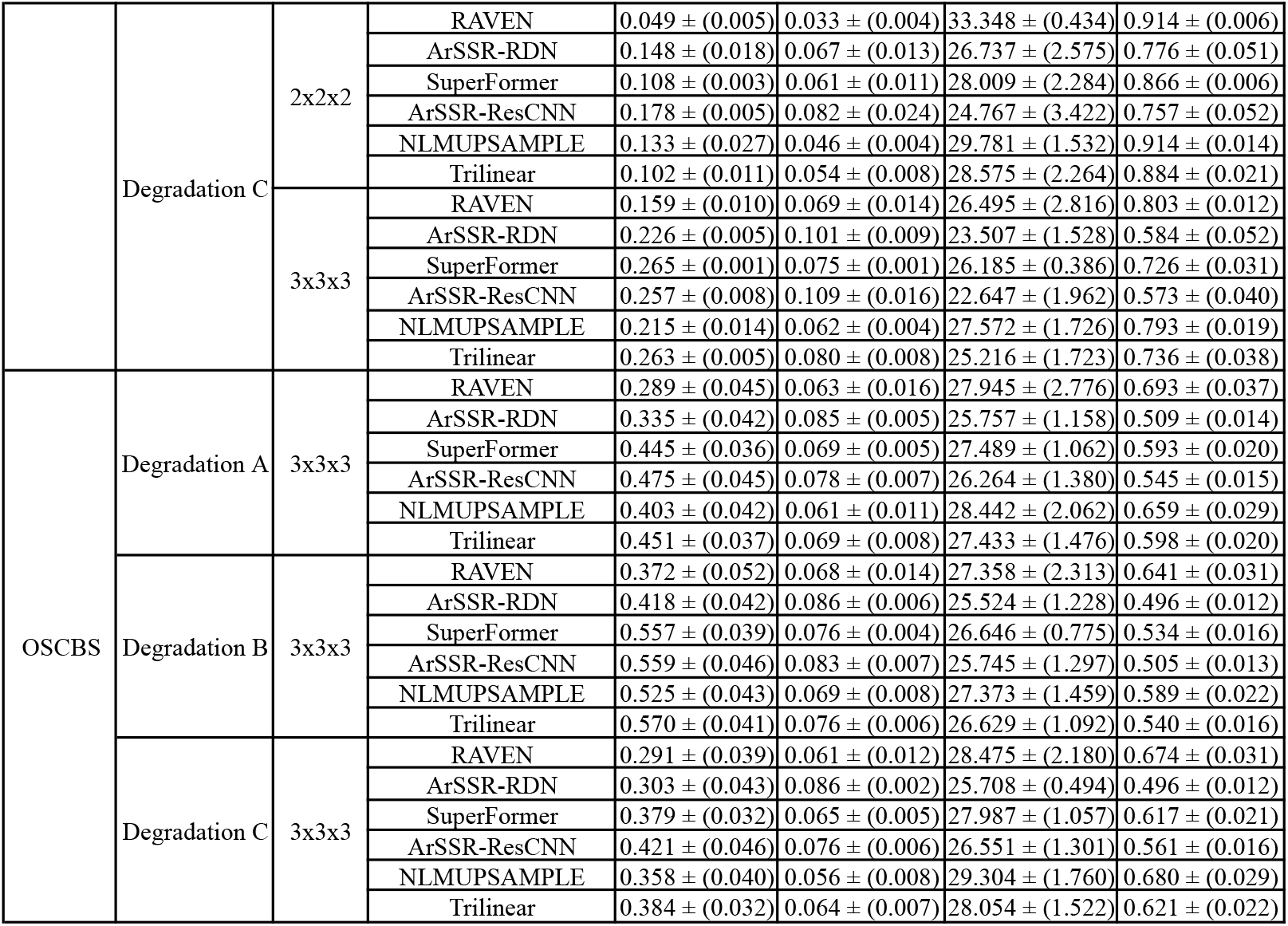
Performance of tested methods per dataset, degradation type and upsampling factor (scale). Metrics reported as “mean ± (standard deviation)”.

**Table S3.** Performance of tested methods per target voxel size, degradation type and upsampling factor (scale). Metrics reported as “mean ± (standard deviation)”.

| Voxel Bin | Degradation | Scale | Upsampling Method | LPIPS ↓ | MAE ↓ | PSNR ↑ | SSIM ↑ |
| --- | --- | --- | --- | --- | --- | --- | --- |
| 0.60-0.90 mm | Degradation A | 2x2x2 | RAVEN | 0.052 ± (0.022) | 0.031 ± (0.018) | 35.842 ± (3.691) | 0.902 ± (0.046) |
| | | | ArSSR-RDN | 0.185 ± (0.046) | 0.050 ± (0.022) | 30.477 ± (3.708) | 0.824 ± (0.056) |
| | | | SuperFormer | 0.149 ± (0.028) | 0.061 ± (0.033) | 29.148 ± (4.025) | 0.825 ± (0.053) |
| | | | ArSSR-ResCNN | 0.236 ± (0.046) | 0.058 ± (0.026) | 29.093 ± (3.794) | 0.782 ± (0.055) |
| | | | NLMUPSAMPLE | 0.172 ± (0.039) | 0.037 ± (0.019) | 34.549 ± (3.677) | 0.876 ± (0.042) |
| | | | Trilinear | 0.132 ± (0.024) | 0.040 ± (0.019) | 34.018 ± (3.625) | 0.852 ± (0.041) |
| | | 3x3x3 | RAVEN | 0.144 ± (0.041) | 0.046 ± (0.026) | 32.085 ± (3.873) | 0.835 ± (0.052) |
| | | | ArSSR-RDN | 0.290 ± (0.053) | 0.075 ± (0.028) | 26.960 ± (3.139) | 0.664 ± (0.062) |
| | | | SuperFormer | 0.352 ± (0.035) | 0.083 ± (0.028) | 26.190 ± (3.171) | 0.663 ± (0.048) |
| | | | ArSSR-ResCNN | 0.347 ± (0.043) | 0.084 ± (0.030) | 25.988 ± (3.346) | 0.624 ± (0.059) |
| | | | NLMUPSAMPLE | 0.302 ± (0.047) | 0.056 ± (0.025) | 30.623 ± (3.407) | 0.752 ± (0.049) |
| | | | Trilinear | 0.344 ± (0.040) | 0.069 ± (0.027) | 28.401 ± (3.141) | 0.683 ± (0.050) |
| | Degradation B | 2x2x2 | RAVEN | 0.059 ± (0.026) | 0.032 ± (0.021) | 35.419 ± (3.955) | 0.910 ± (0.044) |
| | | | ArSSR-RDN | 0.228 ± (0.052) | 0.057 ± (0.026) | 29.248 ± (3.879) | 0.803 ± (0.057) |
| | | | SuperFormer | 0.208 ± (0.033) | 0.066 ± (0.029) | 28.363 ± (3.767) | 0.791 ± (0.048) |
| | | | ArSSR-ResCNN | 0.279 ± (0.049) | 0.063 ± (0.028) | 28.269 ± (3.827) | 0.758 ± (0.055) |
| | | | NLMUPSAMPLE | 0.237 ± (0.046) | 0.045 ± (0.023) | 32.521 ± (3.679) | 0.835 ± (0.045) |
| | | | Trilinear | 0.191 ± (0.035) | 0.049 ± (0.024) | 31.888 ± (3.628) | 0.815 ± (0.045) |
| | | 3x3x3 | RAVEN | 0.206 ± (0.047) | 0.057 ± (0.029) | 29.913 ± (3.656) | 0.775 ± (0.056) |
| | | | ArSSR-RDN | 0.358 ± (0.046) | 0.080 ± (0.031) | 26.786 ± (3.087) | 0.627 ± (0.061) |
| | | | SuperFormer | 0.442 ± (0.035) | 0.096 ± (0.030) | 24.762 ± (3.030) | 0.600 ± (0.050) |
| | | | ArSSR-ResCNN | 0.421 ± (0.040) | 0.087 ± (0.031) | 26.007 ± (3.077) | 0.585 ± (0.059) |
| | | | NLMUPSAMPLE | 0.403 ± (0.046) | 0.070 ± (0.028) | 28.231 ± (3.114) | 0.679 ± (0.050) |
| | | | Trilinear | 0.438 ± (0.041) | 0.080 ± (0.028) | 26.739 ± (2.863) | 0.621 ± (0.051) |
| | Degradation C | 2x2x2 | RAVEN | 0.060 ± (0.030) | 0.031 ± (0.017) | 35.333 ± (3.722) | 0.901 ± (0.044) |
| | | | ArSSR-RDN | 0.187 ± (0.052) | 0.049 ± (0.022) | 30.782 ± (3.480) | 0.821 ± (0.055) |
| | | | SuperFormer | 0.137 ± (0.031) | 0.059 ± (0.025) | 29.288 ± (3.522) | 0.828 ± (0.045) |
| | | | ArSSR-ResCNN | 0.229 ± (0.051) | 0.055 ± (0.024) | 29.535 ± (3.656) | 0.791 ± (0.054) |
| | | | NLMUPSAMPLE | 0.177 ± (0.048) | 0.035 ± (0.019) | 35.025 ± (3.704) | 0.878 ± (0.046) |
| | | | Trilinear | 0.123 ± (0.033) | 0.041 ± (0.021) | 33.508 ± (3.566) | 0.844 ± (0.048) |
| | | 3x3x3 | RAVEN | 0.173 ± (0.045) | 0.047 ± (0.023) | 31.931 ± (3.394) | 0.794 ± (0.054) |
| | | | ArSSR-RDN | 0.280 ± (0.053) | 0.072 ± (0.026) | 27.615 ± (2.837) | 0.655 ± (0.059) |
| | | | SuperFormer | 0.317 ± (0.042) | 0.076 ± (0.026) | 27.073 ± (3.007) | 0.687 ± (0.050) |
| | | | ArSSR-ResCNN | 0.315 ± (0.046) | 0.077 ± (0.028) | 26.841 ± (3.120) | 0.638 ± (0.058) |
| | | | NLMUPSAMPLE | 0.281 ± (0.054) | 0.050 ± (0.022) | 31.417 ± (3.273) | 0.762 ± (0.052) |
| | | | Trilinear | 0.311 ± (0.046) | 0.062 ± (0.026) | 29.447 ± (3.206) | 0.703 ± (0.053) |
| $\leq 0.60$ mm | Degradation A | 2x2x2 | RAVEN | 0.036 ± (0.014) | 0.027 ± (0.002) | 35.472 ± (1.235) | 0.930 ± (0.025) |
| | | | ArSSR-RDN | 0.140 ± (0.036) | 0.062 ± (0.026) | 27.688 ± (5.543) | 0.811 ± (0.077) |
| | | | SuperFormer | 0.151 ± (0.030) | 0.083 ± (0.022) | 25.436 ± (3.644) | 0.830 ± (0.034) |
| | | | ArSSR-ResCNN | 0.195 ± (0.036) | 0.076 ± (0.026) | 25.271 ± (3.323) | 0.770 ± (0.065) |
| | | | NLMUPSAMPLE | 0.134 ± (0.031) | 0.041 ± (0.016) | 33.020 ± (5.119) | 0.905 ± (0.020) |
| | | | Trilinear | 0.127 ± (0.030) | 0.045 ± (0.017) | 32.610 ± (5.470) | 0.882 ± (0.021) |
| | | 3x3x3 | RAVEN | 0.207 ± (0.097) | 0.058 ± (0.021) | 29.091 ± (4.253) | 0.780 ± (0.100) |
| | | | ArSSR-RDN | 0.298 ± (0.061) | 0.088 ± (0.018) | 25.136 ± (2.695) | 0.576 ± (0.084) |
| | | | SuperFormer | 0.401 ± (0.060) | 0.082 ± (0.021) | 25.855 ± (2.740) | 0.642 ± (0.061) |
| | | | ArSSR-ResCNN | 0.397 ± (0.097) | 0.087 ± (0.023) | 25.332 ± (3.079) | 0.578 ± (0.052) |
| | | | NLMUPSAMPLE | 0.333 ± (0.088) | 0.060 ± (0.015) | 29.225 ± (3.569) | 0.729 ± (0.079) |
| | | | Trilinear | 0.396 ± (0.073) | 0.070 ± (0.015) | 27.654 ± (2.833) | 0.663 ± (0.074) |
| | Degradation B | 2x2x2 | RAVEN | $0.042 \pm (0.018)$ | $0.029 \pm (0.006)$ | $34.636 \pm (2.705)$ | $0.936 \pm (0.024)$ |
| | | | ArSSR-RDN | $0.177 \pm (0.045)$ | $0.070 \pm (0.029)$ | $26.529 \pm (5.558)$ | $0.795 \pm (0.065)$ |
| | | | SuperFormer | $0.206 \pm (0.035)$ | $0.086 \pm (0.022)$ | $25.037 \pm (3.417)$ | $0.804 \pm (0.042)$ |
| | | | ArSSR-ResCNN | $0.233 \pm (0.049)$ | $0.072 \pm (0.035)$ | $27.090 \pm (5.267)$ | $0.766 \pm (0.052)$ |
| | | | NLMUPSAMPLE | $0.196 \pm (0.039)$ | $0.050 \pm (0.020)$ | $31.387 \pm (5.338)$ | $0.871 \pm (0.018)$ |
| | | | Trilinear | $0.183 \pm (0.036)$ | $0.054 \pm (0.021)$ | $30.841 \pm (5.266)$ | $0.850 \pm (0.020)$ |
| | | 3x3x3 | RAVEN | $0.286 \pm (0.101)$ | $0.066 \pm (0.022)$ | $28.013 \pm (3.862)$ | $0.725 \pm (0.094)$ |
| | | | ArSSR-RDN | $0.374 \pm (0.069)$ | $0.093 \pm (0.019)$ | $24.548 \pm (2.722)$ | $0.554 \pm (0.069)$ |
| | | | SuperFormer | $0.503 \pm (0.068)$ | $0.088 \pm (0.023)$ | $25.390 \pm (2.755)$ | $0.581 \pm (0.060)$ |
| | | | ArSSR-ResCNN | $0.474 \pm (0.102)$ | $0.089 \pm (0.025)$ | $25.517 \pm (3.179)$ | $0.550 \pm (0.059)$ |
| | | | NLMUPSAMPLE | $0.451 \pm (0.094)$ | $0.071 \pm (0.015)$ | $27.599 \pm (2.847)$ | $0.657 \pm (0.077)$ |
| | | | Trilinear | $0.504 \pm (0.086)$ | $0.079 \pm (0.014)$ | $26.510 \pm (2.318)$ | $0.603 \pm (0.073)$ |
| | Degradation C | 2x2x2 | RAVEN | $0.042 \pm (0.019)$ | $0.027 \pm (0.004)$ | $35.491 \pm (1.595)$ | $0.927 \pm (0.021)$ |
| | | | ArSSR-RDN | $0.140 \pm (0.041)$ | $0.059 \pm (0.023)$ | $28.230 \pm (5.040)$ | $0.808 \pm (0.079)$ |
| | | | SuperFormer | $0.137 \pm (0.030)$ | $0.084 \pm (0.023)$ | $25.383 \pm (3.300)$ | $0.831 \pm (0.037)$ |
| | | | ArSSR-ResCNN | $0.186 \pm (0.035)$ | $0.073 \pm (0.024)$ | $25.645 \pm (2.981)$ | $0.776 \pm (0.070)$ |
| | | | NLMUPSAMPLE | $0.127 \pm (0.035)$ | $0.036 \pm (0.012)$ | $34.380 \pm (5.174)$ | $0.910 \pm (0.025)$ |
| | | | Trilinear | $0.113 \pm (0.031)$ | $0.044 \pm (0.014)$ | $32.689 \pm (5.114)$ | $0.878 \pm (0.025)$ |
| | | 3x3x3 | RAVEN | $0.225 \pm (0.080)$ | $0.056 \pm (0.018)$ | $29.626 \pm (3.912)$ | $0.751 \pm (0.087)$ |
| | | | ArSSR-RDN | $0.279 \pm (0.054)$ | $0.086 \pm (0.013)$ | $25.472 \pm (1.838)$ | $0.562 \pm (0.086)$ |
| | | | SuperFormer | $0.350 \pm (0.051)$ | $0.080 \pm (0.023)$ | $26.197 \pm (2.821)$ | $0.663 \pm (0.057)$ |
| | | | ArSSR-ResCNN | $0.355 \pm (0.084)$ | $0.084 \pm (0.019)$ | $25.567 \pm (2.399)$ | $0.589 \pm (0.055)$ |
| | | | NLMUPSAMPLE | $0.297 \pm (0.081)$ | $0.053 \pm (0.010)$ | $30.154 \pm (2.932)$ | $0.743 \pm (0.072)$ |
| | | | Trilinear | $0.343 \pm (0.062)$ | $0.064 \pm (0.013)$ | $28.501 \pm (2.920)$ | $0.686 \pm (0.072)$ |
| > 0.90 mm | Degradation A | 2x2x2 | RAVEN | $0.052 \pm (0.010)$ | $0.028 \pm (0.005)$ | $35.905 \pm (1.757)$ | $0.895 \pm (0.025)$ |
| | | | ArSSR-RDN | $0.193 \pm (0.063)$ | $0.042 \pm (0.039)$ | $32.534 \pm (2.673)$ | $0.825 \pm (0.069)$ |
| | | | SuperFormer | $0.143 \pm (0.015)$ | $0.051 \pm (0.015)$ | $29.884 \pm (2.554)$ | $0.828 \pm (0.029)$ |
| | | | ArSSR-ResCNN | $0.225 \pm (0.025)$ | $0.045 \pm (0.007)$ | $31.229 \pm (1.491)$ | $0.783 \pm (0.031)$ |
| | | | NLMUPSAMPLE | $0.195 \pm (0.034)$ | $0.035 \pm (0.017)$ | $33.904 \pm (1.661)$ | $0.867 \pm (0.036)$ |
| | | | Trilinear | $0.125 \pm (0.012)$ | $0.037 \pm (0.005)$ | $33.456 \pm (1.343)$ | $0.848 \pm (0.025)$ |
| | | 3x3x3 | RAVEN | $0.157 \pm (0.023)$ | $0.040 \pm (0.006)$ | $32.324 \pm (1.281)$ | $0.826 \pm (0.026)$ |
| | | | ArSSR-RDN | $0.265 \pm (0.020)$ | $0.067 \pm (0.007)$ | $27.728 \pm (1.048)$ | $0.662 \pm (0.032)$ |
| | | | SuperFormer | $0.311 \pm (0.020)$ | $0.074 \pm (0.014)$ | $26.457 \pm (1.708)$ | $0.661 \pm (0.038)$ |
| | | | ArSSR-ResCNN | $0.309 \pm (0.021)$ | $0.073 \pm (0.007)$ | $26.990 \pm (1.034)$ | $0.613 \pm (0.034)$ |
| | | | NLMUPSAMPLE | $0.276 \pm (0.023)$ | $0.052 \pm (0.006)$ | $29.958 \pm (1.076)$ | $0.747 \pm (0.030)$ |
| | | | Trilinear | $0.307 \pm (0.020)$ | $0.065 \pm (0.006)$ | $27.789 \pm (0.872)$ | $0.673 \pm (0.030)$ |
| | Degradation B | 2x2x2 | RAVEN | $0.063 \pm (0.013)$ | $0.027 \pm (0.005)$ | $36.090 \pm (1.696)$ | $0.905 \pm (0.023)$ |
| | | | ArSSR-RDN | $0.222 \pm (0.025)$ | $0.045 \pm (0.006)$ | $31.095 \pm (1.287)$ | $0.811 \pm (0.032)$ |
| | | | SuperFormer | $0.190 \pm (0.017)$ | $0.055 \pm (0.014)$ | $29.102 \pm (2.190)$ | $0.796 \pm (0.032)$ |
| | | | ArSSR-ResCNN | $0.256 \pm (0.025)$ | $0.052 \pm (0.007)$ | $29.992 \pm (1.246)$ | $0.758 \pm (0.033)$ |
| | | | NLMUPSAMPLE | $0.247 \pm (0.075)$ | $0.046 \pm (0.049)$ | $31.776 \pm (2.590)$ | $0.826 \pm (0.075)$ |
| | | | Trilinear | $0.173 \pm (0.015)$ | $0.045 \pm (0.005)$ | $31.349 \pm (1.155)$ | $0.812 \pm (0.027)$ |
| | | 3x3x3 | RAVEN | $0.205 \pm (0.021)$ | $0.050 \pm (0.006)$ | $30.017 \pm (1.071)$ | $0.763 \pm (0.029)$ |
| | | | ArSSR-RDN | $0.318 \pm (0.018)$ | $0.076 \pm (0.006)$ | $26.510 \pm (0.806)$ | $0.618 \pm (0.032)$ |
| | | | SuperFormer | $0.387 \pm (0.021)$ | $0.084 \pm (0.011)$ | $25.225 \pm (1.219)$ | $0.589 \pm (0.036)$ |
| | | | ArSSR-ResCNN | $0.381 \pm (0.018)$ | $0.082 \pm (0.007)$ | $25.797 \pm (0.840)$ | $0.563 \pm (0.033)$ |
| | | | NLMUPSAMPLE | $0.356 \pm (0.021)$ | $0.067 \pm (0.006)$ | $27.533 \pm (0.851)$ | $0.664 \pm (0.031)$ |
| | | | Trilinear | $0.387 \pm (0.021)$ | $0.077 \pm (0.006)$ | $26.182 \pm (0.750)$ | $0.600 \pm (0.031)$ |
| | Degradation C | 2x2x2 | RAVEN | $0.067 \pm (0.013)$ | $0.027 \pm (0.005)$ | $35.793 \pm (1.615)$ | $0.901 \pm (0.022)$ |
| | | | ArSSR-RDN | $0.196 \pm (0.028)$ | $0.039 \pm (0.007)$ | $32.729 \pm (1.595)$ | $0.830 \pm (0.031)$ |
| | | | SuperFormer | $0.136 \pm (0.016)$ | $0.051 \pm (0.014)$ | $29.836 \pm (2.399)$ | $0.832 \pm (0.028)$ |
| | | | ArSSR-ResCNN | $0.223 \pm (0.026)$ | $0.043 \pm (0.007)$ | $31.573 \pm (1.491)$ | $0.794 \pm (0.030)$ |
| | | | NLMUPSAMPLE | $0.205 \pm (0.027)$ | $0.032 \pm (0.005)$ | $34.372 \pm (1.526)$ | $0.875 \pm (0.026)$ |
| | | | Trilinear | $0.119 \pm (0.013)$ | $0.037 \pm (0.005)$ | $33.103 \pm (1.338)$ | $0.846 \pm (0.026)$ |
| | | 3x3x3 | RAVEN | $0.184 \pm (0.024)$ | $0.041 \pm (0.006)$ | $32.254 \pm (1.303)$ | $0.803 \pm (0.027)$ |
| | | | ArSSR-RDN | $0.258 \pm (0.022)$ | $0.064 \pm (0.008)$ | $28.267 \pm (1.263)$ | $0.665 \pm (0.030)$ |
| | | | SuperFormer | $0.291 \pm (0.020)$ | $0.067 \pm (0.013)$ | $27.391 \pm (1.759)$ | $0.696 \pm (0.033)$ |
| | | | ArSSR-ResCNN | $0.283 \pm (0.022)$ | $0.066 \pm (0.009)$ | $27.942 \pm (1.319)$ | $0.638 \pm (0.033)$ |
| | | | NLMUPSAMPLE | $0.269 \pm (0.026)$ | $0.046 \pm (0.006)$ | $31.082 \pm (1.149)$ | $0.771 \pm (0.028)$ |
| | | | Trilinear | $0.290 \pm (0.021)$ | $0.058 \pm (0.005)$ | $28.988 \pm (0.940)$ | $0.708 \pm (0.029)$ |

**Figure S1.**
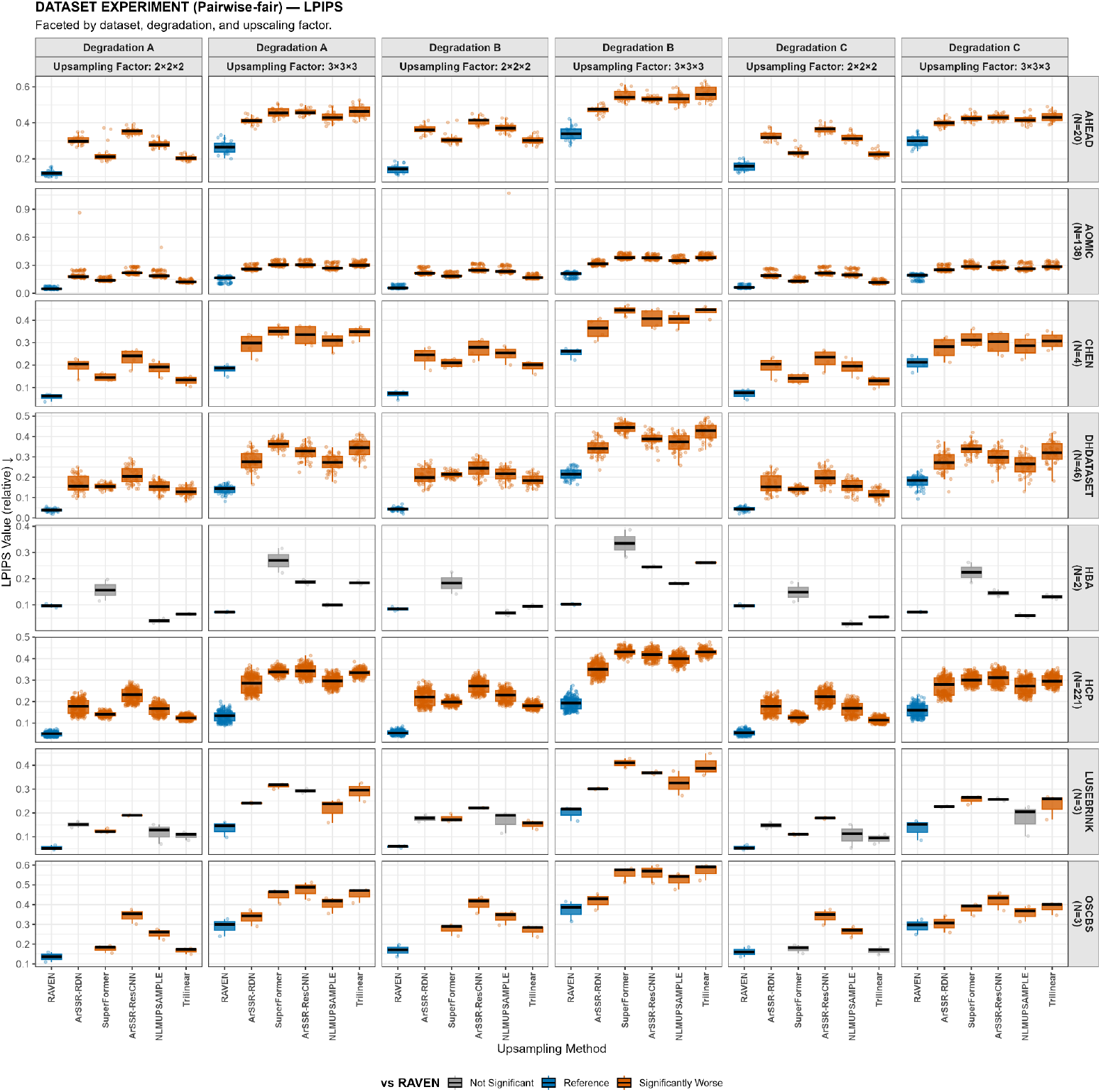
LPIPS (lower is better) benchmark results for super-resolution tasks across datasets and upsampling factors. The color of NLMUPSAMPLE and Trilinear boxplots indicate if no significant differences were found with RAVEN (gray), or if the given method was significantly better (green) or worse (orange) than RAVEN.

**Figure S2.**
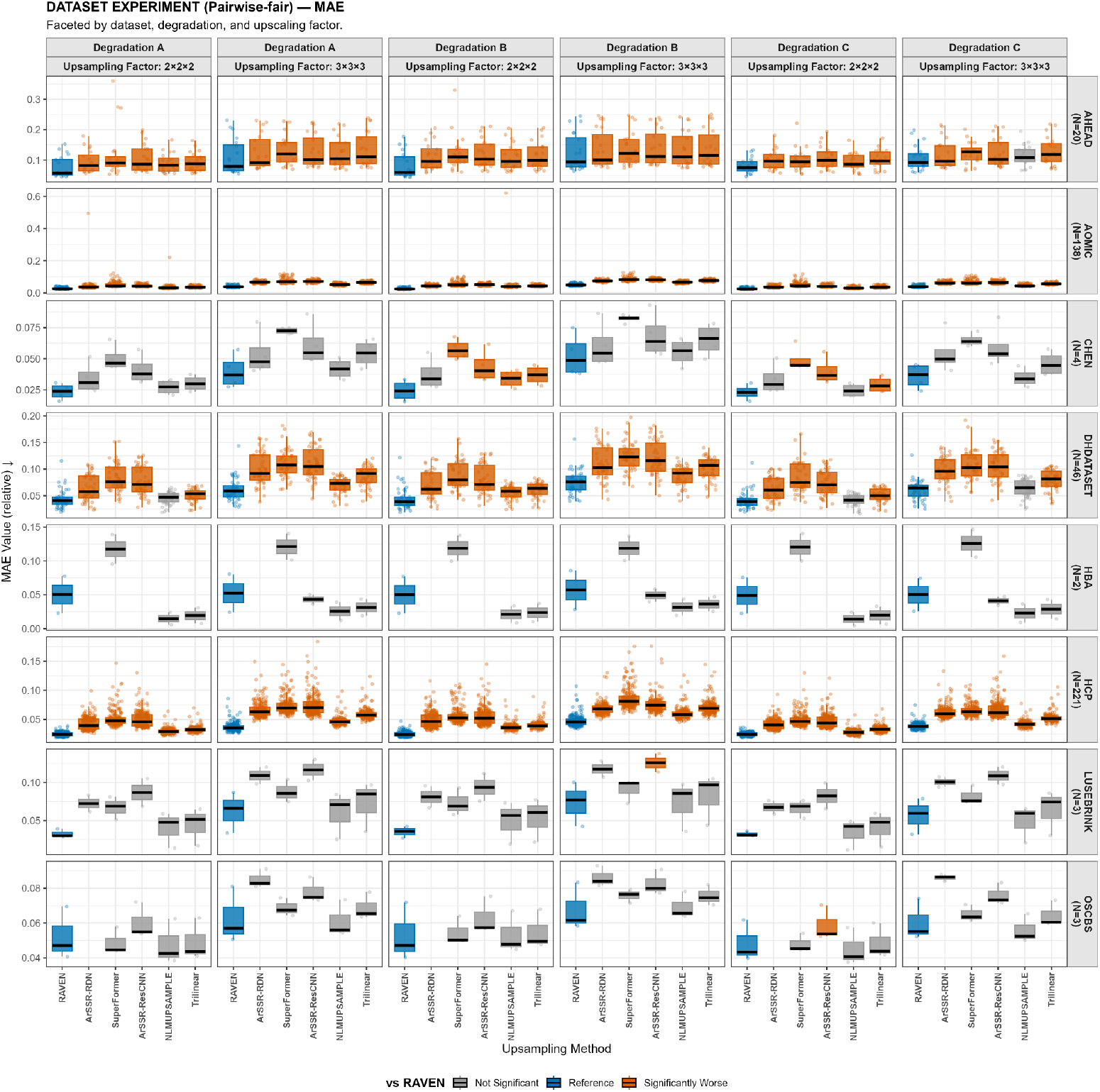
MAE (lower is better) benchmark results for super-resolution tasks across datasets and upsampling factors. The color of NLMUPSAMPLE and Trilinear boxplots indicate if no significant differences were found with RAVEN (gray), or if the given method was significantly better (green) or worse (orange) than RAVEN.

**Figure S3.**
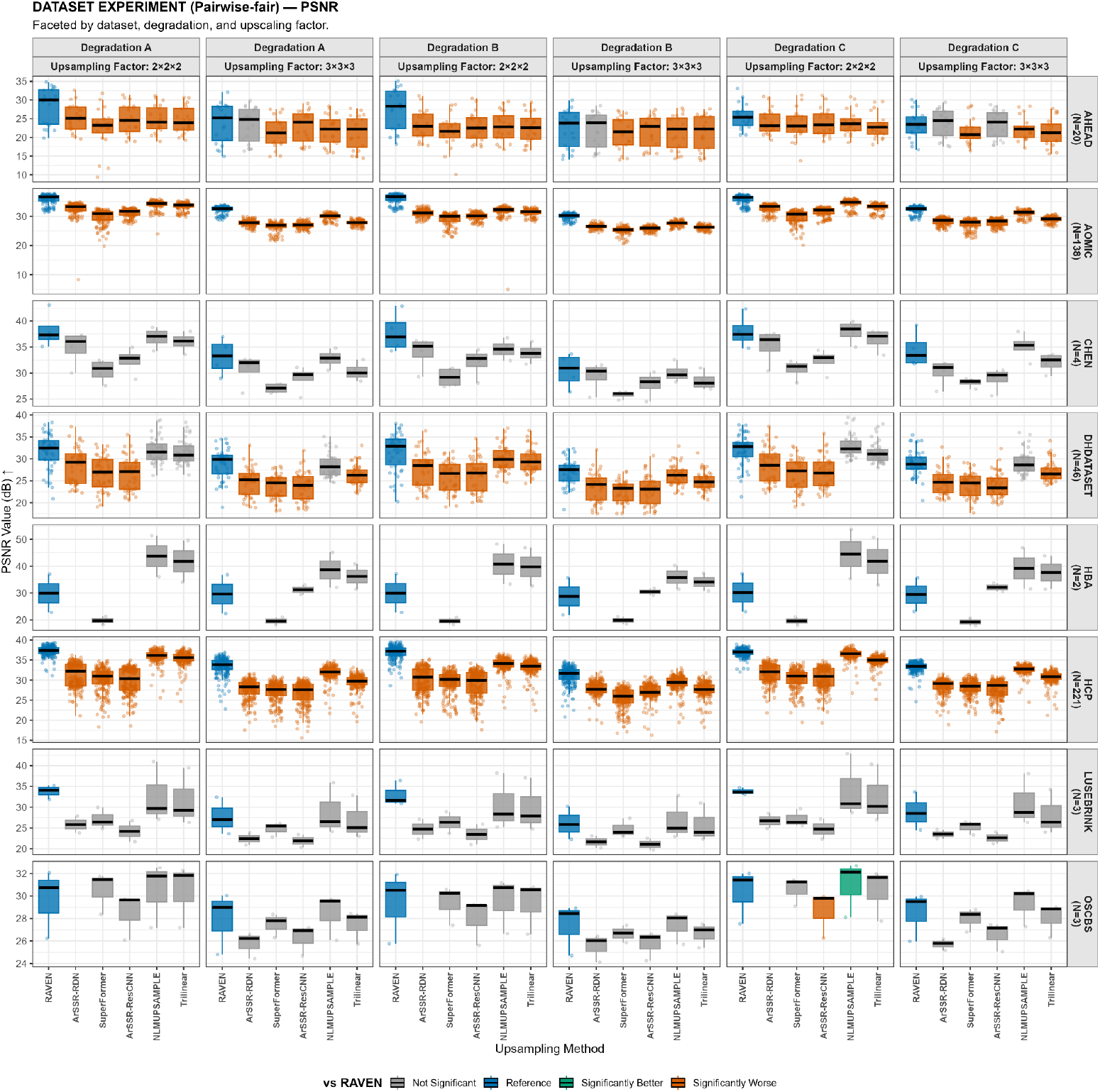
PSNR (higher is better, in dB) benchmark results for super-resolution tasks across datasets and upsampling factors. The color of NLMUPSAMPLE and Trilinear boxplots indicate if no significant differences were found with RAVEN (gray), or if the given method was significantly better (green) or worse (orange) than RAVEN.

**Figure S4.**
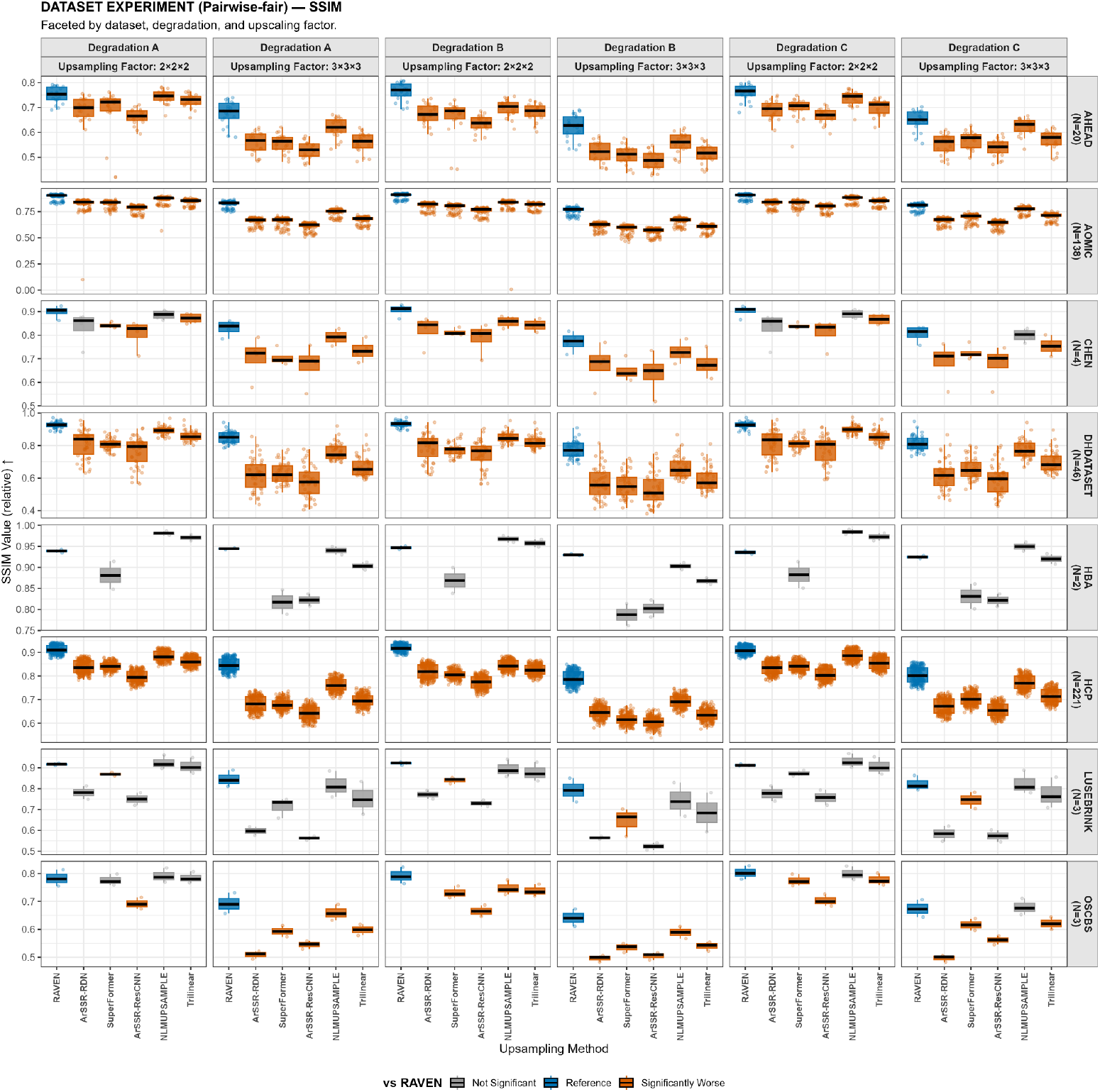
SSIM (higher is better) benchmark results for super-resolution tasks across datasets and upsampling factors. The color of NLMUPSAMPLE and Trilinear boxplots indicate if no significant differences were found with RAVEN (gray), or if the given method was significantly better (green) or worse (orange) than RAVEN.

**Figure S5.**
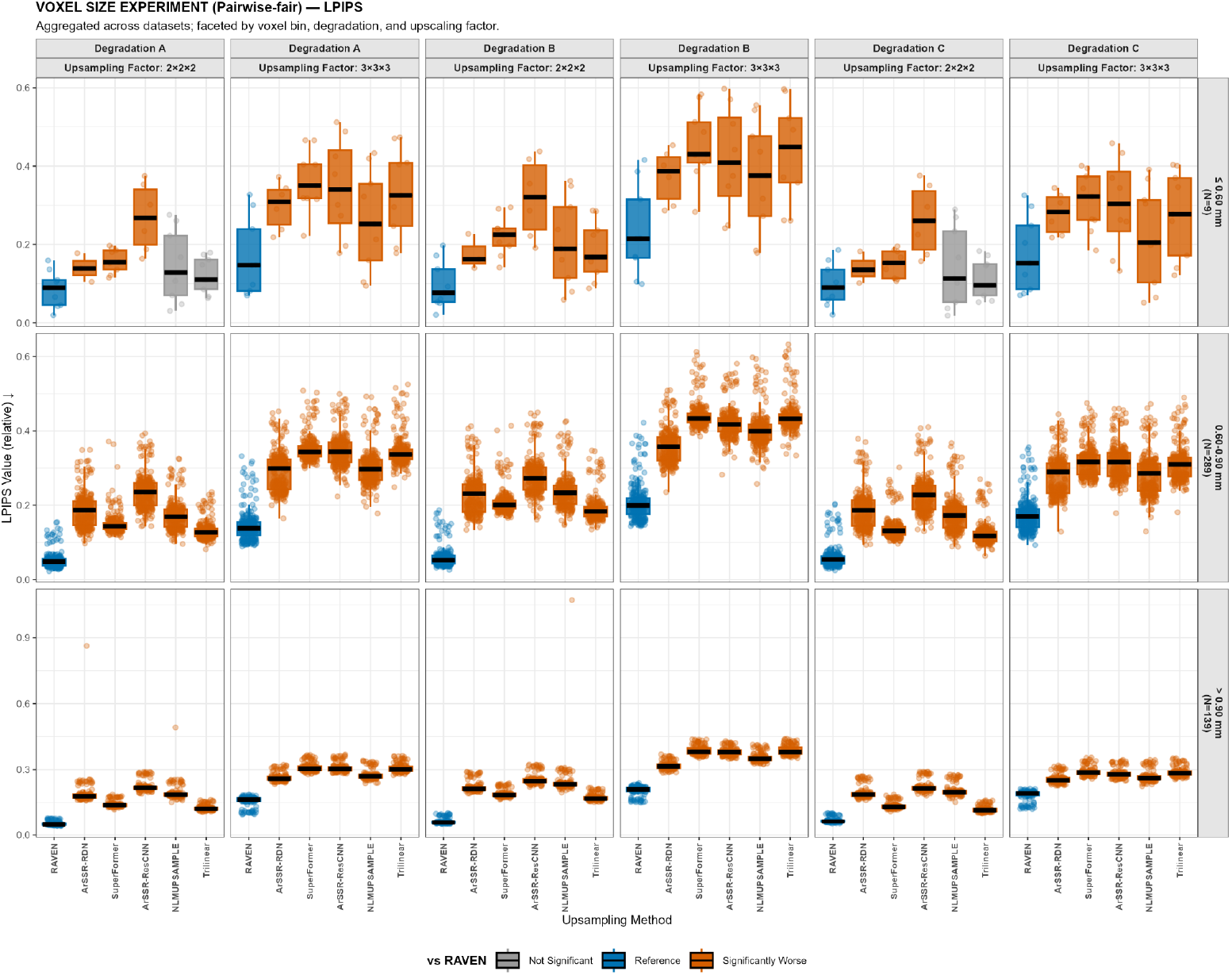
LPIPS (lower is better) benchmark results for super-resolution tasks across different target voxel sizes (row names on the right) and upsampling factors. The color of NLMUPSAMPLE and Trilinear boxplots indicate if no significant differences were found with RAVEN (gray), or if the given method was significantly better (green) or worse (orange) than RAVEN.

**Figure S6.**
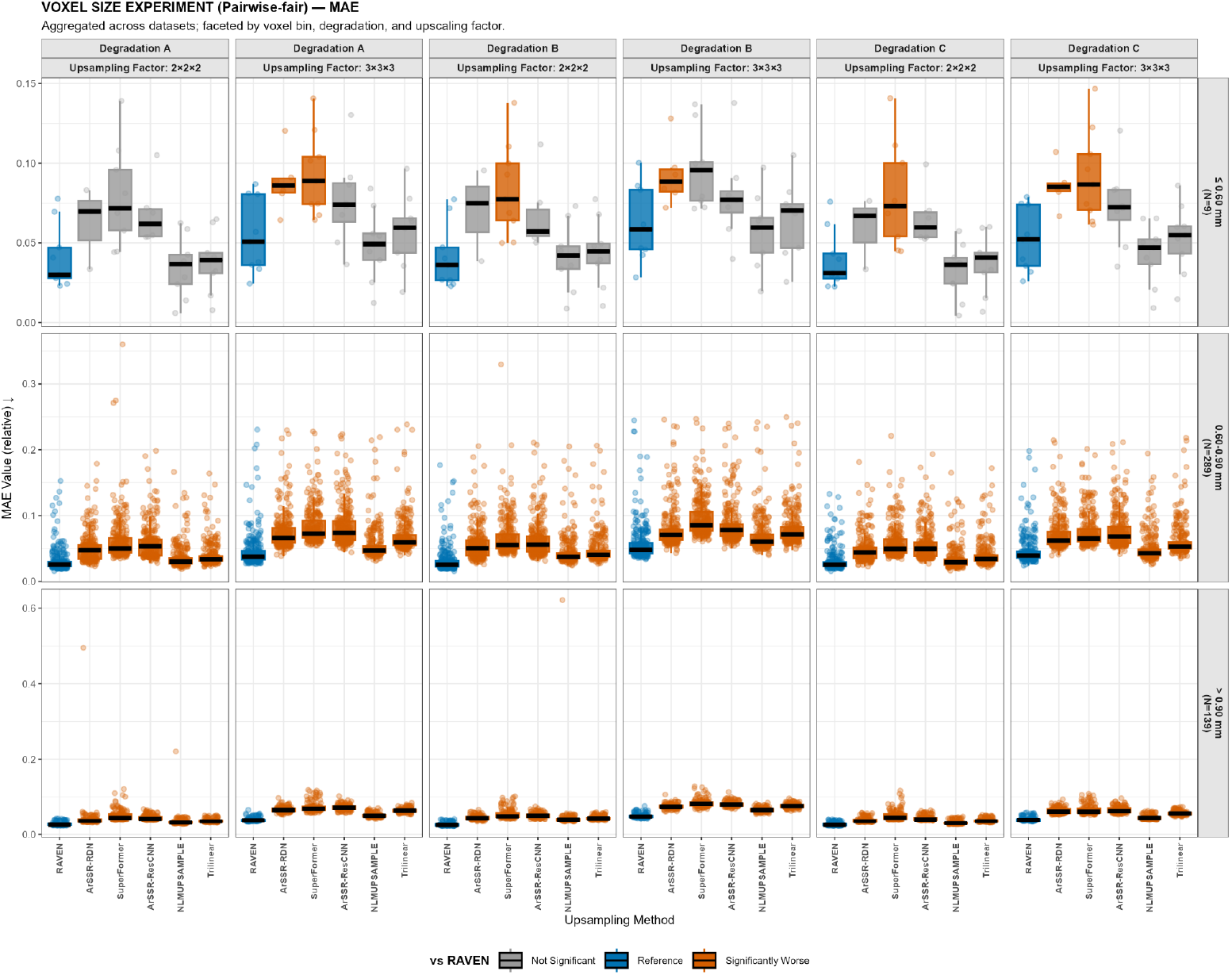
MAE (lower is better) benchmark results for super-resolution tasks across different target voxel sizes (row names on the right) and upsampling factors. The color of NLMUPSAMPLE and Trilinear boxplots indicate if no significant differences were found with RAVEN (gray), or if the given method was significantly better (green) or worse (orange) than RAVEN.

**Figure S7.**
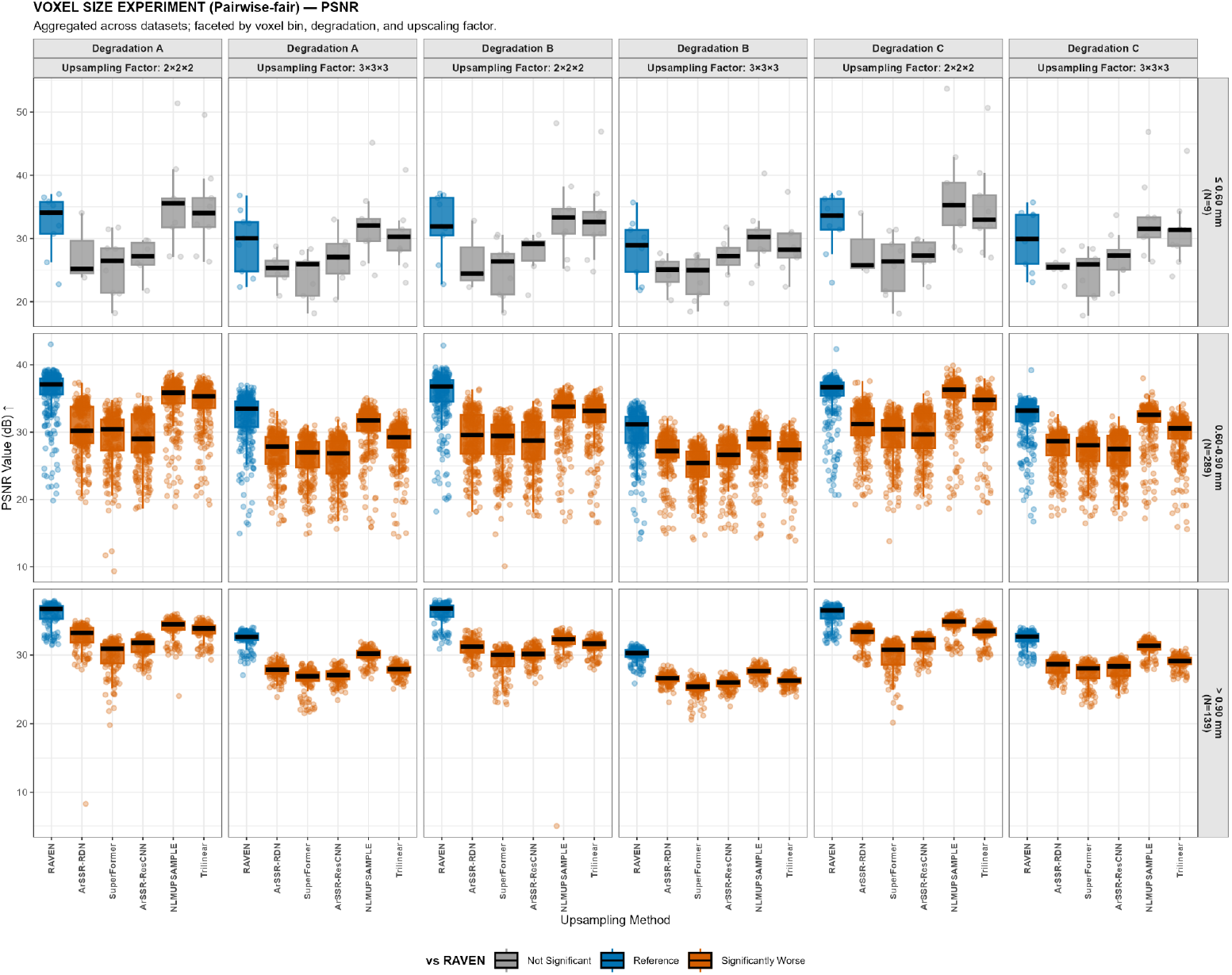
PSNR (higher is better, in dB) benchmark results for super-resolution tasks across different target voxel sizes (row names on the right) and upsampling factors. The color of NLMUPSAMPLE and Trilinear boxplots indicate if no significant differences were found with RAVEN (gray), or if the given method was significantly better (green) or worse (orange) than RAVEN.

**Figure S8.**
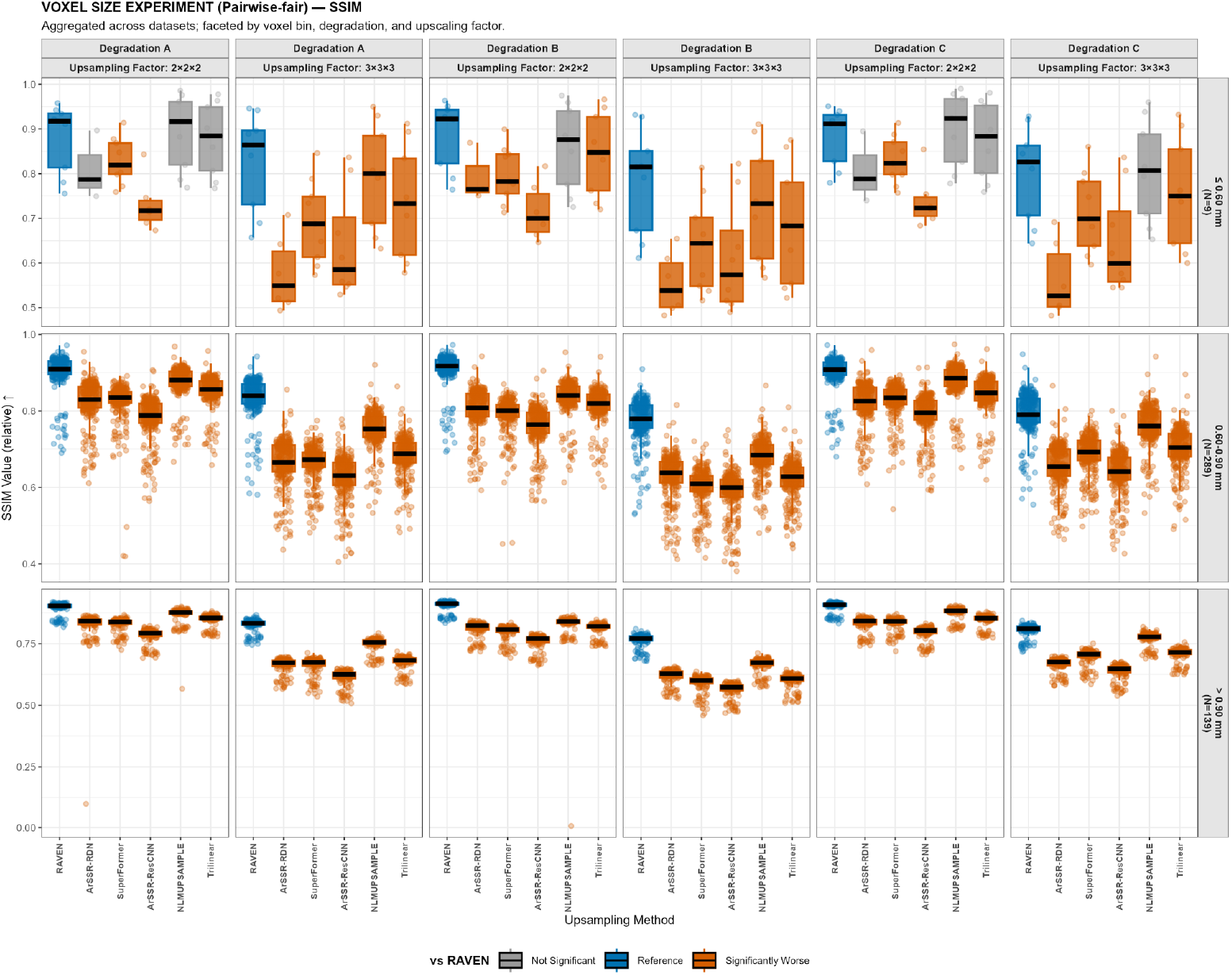
SSIM (higher is better) benchmark results for super-resolution tasks across different target voxel sizes (row names on the right) and upsampling factors. The color of NLMUPSAMPLE and Trilinear boxplots indicate if no significant differences were found with RAVEN (gray), or if the given method was significantly better (green) or worse (orange) than RAVEN.

**Figure S9.**
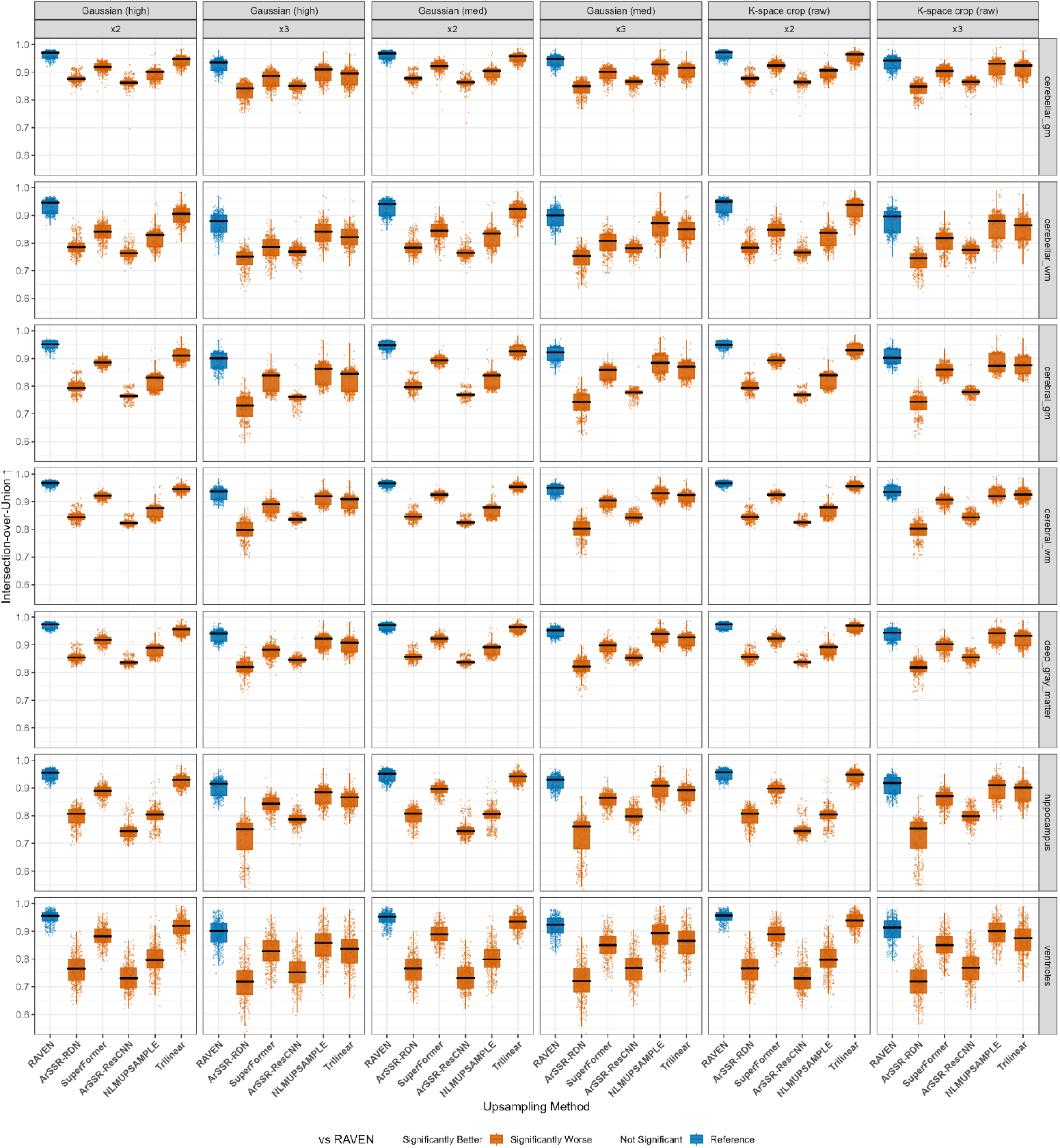
IOU (higher is better) for 7 regions of interest generated by SynthSeg after resampling them to the input’s native space. Results separated for different degradation methods and upsampling factors. Blue is the reference method (RAVEN), and orange depicts statistically worse results vs the reference method after FDR correction.

**Figure S10.**
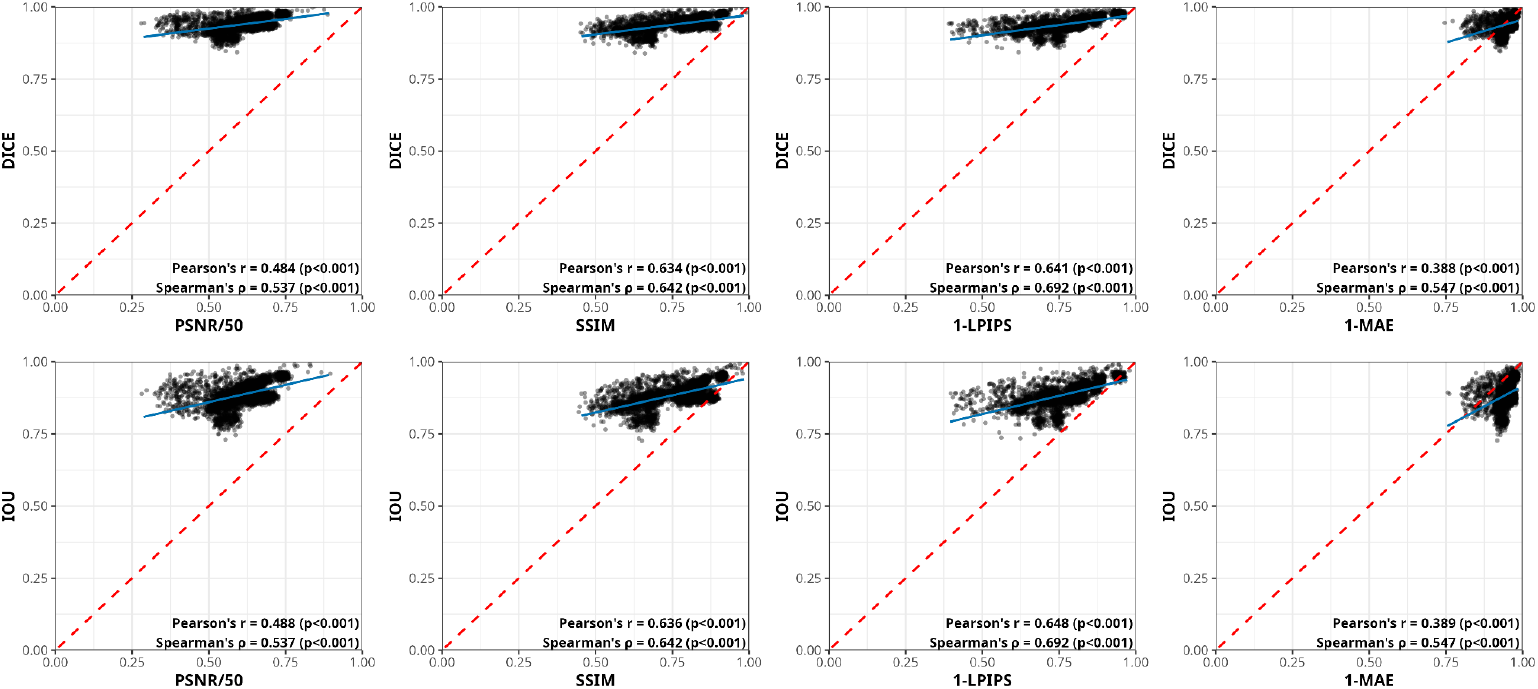
Correlation between image similarity metrics (PSNR, SSIM, LPIPS, and MAE), and segmentation agreement metrics (IOU and DICE) for bilateral hippocampal labels generated by SynthSeg. The correlation between 1-LPIPS was the highest compared to 1-MAE, SSIM, and PSNR/50 (PSNR divided by 50 for visualization purposes).

## Notes

https://github.com/waadgo/raven

### Competing Interest Statement

The authors have declared no competing interest.

## References

Adame-Gonzalez, W., Brzezinski-Rittner, A., Chakravarty, M. M., Farivar, R., & Dadar, M. (2023). FONDUE: Robust resolution-invariant denoising of MR Images using Nested UNets [Preprint]. Neuroscience. 10.1101/2023.06.04.543602

Alkemade, A., Mulder, M. J., Groot, J. M., Isaacs, B. R., Van Berendonk, N., Lute, N., Isherwood, S. J., Bazin, P.-L., & Forstmann, B. U. (2020). The Amsterdam Ultra-high field adult lifespan database (AHEAD): A freely available multimodal 7 Tesla submillimeter magnetic resonance imaging database. NeuroImage, 221, 117200. 10.1016/j.neuroimage.2020.117200

Avants, B., Tustison, N. J., McMillan, C. T., Gosselin, T., Gunn, R., & Hesterman, J. (2023). Concurrent 3D super resolution on intensity and segmentation maps improves detection of structural effects in neurodegenerative disease. Radiology and Imaging. 10.1101/2023.02.02.23285376

Benjamini, Y., & Hochberg, Y. (1995). Controlling the False Discovery Rate: A Practical and Powerful Approach to Multiple Testing. Journal of the Royal Statistical Society Series B: Statistical Methodology, 57(1), 289–300. 10.1111/j.2517-6161.1995.tb02031.x

Billot, B., Greve, D. N., Puonti, O., Thielscher, A., Van Leemput, K., Fischl, B., Dalca, A. V., & Iglesias, J. E. (2023). SynthSeg: Segmentation of brain MRI scans of any contrast and resolution without retraining. Medical Image Analysis, 86, 102789. 10.1016/j.media.2023.102789

Chen, X., Qu, L., Xie, Y., Ahmad, S., & Yap, P.-T. (2023). A paired dataset of T1-and T2-weighted MRI at 3 Tesla and 7 Tesla. Scientific Data, 10(1), 489. 10.1038/s41597-023-02400-y

Chen, Y., Xie, Y., Zhou, Z., Shi, F., Christodoulou, A. G., & Li, D. (2018). Brain MRI super resolution using 3D deep densely connected neural networks. 2018 IEEE 15th International Symposium on Biomedical Imaging (ISBI 2018), 739–742. 10.1109/ISBI.2018.8363679

Dadar, M., Moqadam, R., Metz, A., Chadwick, K., & Zeighami, Y. (n.d.). PELICAN: a Longitudinal Image Processing Pipeline for Analyzing Structural Magnetic Resonance Images in Aging and Neurodegenerative Disease Populations.

Dadar, M., Sanches, L., Fouquet, J. P., Moqadam, R., Alasmar, Z., Ruth Leppert, I., Mirault, D., Maranzano, J., Mechawar, N., Chakravarty, M., & Zeighami, Y. (2024). The Douglas-Bell Canada Brain Bank Post-mortem Brain Imaging Protocol. Aperture Neuro, 4. 10.52294/001c.123347

Demir, U., & Unal, G. (2018). Patch-Based Image Inpainting with Generative Adversarial Networks (No. 1803.07422). arXiv. 10.48550/arXiv.1803.07422

Dice, L. R. (1945). Measures of the Amount of Ecologic Association Between Species. Ecology, 26(3), 297–302. 10.2307/1932409

Esser, P., Rombach, R., & Ommer, B. (2021). Taming Transformers for High-Resolution Image Synthesis (No. 2012.09841). arXiv. 10.48550/arXiv.2012.09841

Fischer, J. S., Gui, M., Ma, P., Stracke, N., Baumann, S. A., & Ommer, B. (2024). Boosting Latent Diffusion with Flow Matching (No. 2312.07360). arXiv. http://arxiv.org/abs/2312.07360

Forigua, C., Escobar, M., & Arbelaez, P. (2022). SuperFormer: Volumetric Transformer Architectures for MRI Super-Resolution. In C. Zhao, D. Svoboda, J. M. Wolterink, & M. Escobar (Eds.), Simulation and Synthesis in Medical Imaging (Vol. 13570, pp. 132–141). Springer International Publishing. 10.1007/978-3-031-16980-9_13

Haris, M., Shakhnarovich, G., & Ukita, N. (2018). Deep Back-Projection Networks For Super-Resolution (No. 1803.02735). arXiv. 10.48550/arXiv.1803.02735

Henschel, L., Kügler, D., & Reuter, M. (2022). FastSurferVINN: Building resolution-independence into deep learning segmentation methods—A solution for HighRes brain MRI. NeuroImage, 251, 118933. 10.1016/j.neuroimage.2022.118933

Iglesias, J. E., Billot, B., Balbastre, Y., Tabari, A., Conklin, J., Gilberto González, R., Alexander, D. C., Golland, P., Edlow, B. L., & Fischl, B. (2021). Joint super-resolution and synthesis of 1 mm isotropic MP-RAGE volumes from clinical MRI exams with scans of different orientation, resolution and contrast. NeuroImage, 237, 118206. 10.1016/j.neuroimage.2021.118206

Jaccard, P. (1901). Étude comparative de la distribution florale dans une portion des Alpes et du Jura [Text/html,application/pdf,text/html]. 10.5169/SEALS-266450

Jo, Y., Yang, S., & Kim, S. J. (2020). Investigating Loss Functions for Extreme Super-Resolution. 2020 IEEE/CVF Conference on Computer Vision and Pattern Recognition Workshops (CVPRW), 1705–1712. 10.1109/CVPRW50498.2020.00220

Lin, J., Miao, Q., Surawech, C., Raman, S. S., Zhao, K., Wu, H. H., & Sung, K. (2023). High-Resolution 3D MRI With Deep Generative Networks via Novel Slice-Profile Transformation Super-Resolution. IEEE Access, 11, 95022–95036. 10.1109/ACCESS.2023.3307577

Lüsebrink, F., Mattern, H., Yakupov, R., Acosta-Cabronero, J., Ashtarayeh, M., Oeltze-Jafra, S., & Speck, O. (2021). Comprehensive ultrahigh resolution whole brain in vivo MRI dataset as a human phantom. Scientific Data, 8(1), 138. 10.1038/s41597-021-00923-w

Lüsebrink, F., Sciarra, A., Mattern, H., Yakupov, R., & Speck, O. (2017). T1-weighted in vivo human whole brain MRI dataset with an ultrahigh isotropic resolution of 250 μm. Scientific Data, 4(1), 170032. 10.1038/sdata.2017.32

Lüsebrink, F., Sciarra, A., Mattern, H., Yakupov, R., & Speck, O. (2018). Data from: T1-weighted in vivo human whole brain MRI dataset with an ultrahigh isotropic resolution of 250 μm (Version 1, p. 20692776990 bytes) [Dataset]. Dryad. 10.5061/DRYAD.38S74

Lyu, Q., You, C., Shan, H., & Wang, G. (2018). Super-resolution MRI through Deep Learning (No. 1810.06776). arXiv. 10.48550/arXiv.1810.06776

Manjón, J. V., Coupé, P., Buades, A., Fonov, V., Louis Collins, D., & Robles, M. (2010). Non-local MRI upsampling. Medical Image Analysis, 14(6), 784–792. 10.1016/j.media.2010.05.010

Mueller, S. G., Weiner, M. W., Thal, L. J., Petersen, R. C., Jack, C. R., Jagust, W., Trojanowski, J. Q., Toga, A. W., & Beckett, L. (2005). Ways toward an early diagnosis in Alzheimer’s disease: The Alzheimer’s Disease Neuroimaging Initiative (ADNI). Alzheimer’s & Dementia, 1(1), 55–66. 10.1016/j.jalz.2005.06.003

Rombach, R., Blattmann, A., Lorenz, D., Esser, P., & Ommer, B. (2022). High-Resolution Image Synthesis with Latent Diffusion Models (No. 2112.10752). arXiv. http://arxiv.org/abs/2112.10752

Schira, M. M., Isherwood, Z. J., Kassem, M. S., Barth, M., Shaw, T. B., Roberts, M. M., & Paxinos, G. (2023). HumanBrainAtlas: An in vivo MRI dataset for detailed segmentations. Brain Structure and Function, 228(8), 1849–1863. 10.1007/s00429-023-02653-8

Shafiee, N., Fonov, V., Dadar, M., Spreng, R. N., & Collins, D. L. (2024). Degeneration in Nucleus basalis of Meynert signals earliest stage of Alzheimer’s disease progression. Neurobiology of Aging, 139, 54–63. 10.1016/j.neurobiolaging.2024.03.003

Shi, W., Caballero, J., Huszar, F., Totz, J., Aitken, A. P., Bishop, R., Rueckert, D., & Wang, Z. (2016). Real-Time Single Image and Video Super-Resolution Using an Efficient Sub-Pixel Convolutional Neural Network. 2016 IEEE Conference on Computer Vision and Pattern Recognition (CVPR), 1874–1883. 10.1109/CVPR.2016.207

Snoek, L., Van Der Miesen, M. M., Beemsterboer, T., Van Der Leij, A., Eigenhuis, A., & Steven Scholte, H. (2021). The Amsterdam Open MRI Collection, a set of multimodal MRI datasets for individual difference analyses. Scientific Data, 8(1), 85. 10.1038/s41597-021-00870-6

Tardif, C. L., Schäfer, A., Trampel, R., Villringer, A., Turner, R., & Bazin, P.-L. (2016). Open Science CBS Neuroimaging Repository: Sharing ultra-high-field MR images of the brain. NeuroImage, 124, 1143–1148. 10.1016/j.neuroimage.2015.08.042

Tustison, N. J., Cook, P. A., Holbrook, A. J., Johnson, H. J., Muschelli, J., Devenyi, G. A., Duda, J. T., Das, S. R., Cullen, N. C., Gillen, D. L., Yassa, M. A., Stone, J. R., Gee, J. C., & Avants, B. B. (2021). The ANTsX ecosystem for quantitative biological and medical imaging. Scientific Reports, 11(1), 9068. 10.1038/s41598-021-87564-6

Umer, R. M., Foresti, G. L., & Micheloni, C. (2020). Deep Generative Adversarial Residual Convolutional Networks for Real-World Super-Resolution (No. 2005.00953). arXiv. http://arxiv.org/abs/2005.00953

Umer, R. M., & Micheloni, C. (2020). Deep Cyclic Generative Adversarial Residual Convolutional Networks for Real Image Super-Resolution (No. 2009.03693). arXiv. http://arxiv.org/abs/2009.03693

Van Essen, D. C., Ugurbil, K., Auerbach, E., Barch, D., Behrens, T. E. J., Bucholz, R., Chang, A., Chen, L., Corbetta, M., Curtiss, S. W., Della Penna, S., Feinberg, D., Glasser, M. F., Harel, N., Heath, A. C., Larson-Prior, L., Marcus, D., Michalareas, G., Moeller, S., … Yacoub, E. (2012). The Human Connectome Project: A data acquisition perspective. NeuroImage, 62(4), 2222–2231. 10.1016/j.neuroimage.2012.02.018

Wang, J., Levman, J., Pinaya, W. H. L., Tudosiu, P.-D., Cardoso, M. J., & Marinescu, R. (2023). InverseSR: 3D Brain MRI Super-Resolution Using a Latent Diffusion Model (No. 2308.12465). arXiv. http://arxiv.org/abs/2308.12465

Wang, X., Yu, K., Wu, S., Gu, J., Liu, Y., Dong, C., Loy, C. C., Qiao, Y., & Tang, X. (2018). ESRGAN: Enhanced Super-Resolution Generative Adversarial Networks (No. 1809.00219). arXiv. http://arxiv.org/abs/1809.00219

Wu, Q., Li, Y., Sun, Y., Zhou, Y., Wei, H., Yu, J., & Zhang, Y. (2023). An Arbitrary Scale Super-Resolution Approach for 3D MR Images via Implicit Neural Representation. IEEE Journal of Biomedical and Health Informatics, 27(2), 1004–1015. 10.1109/JBHI.2022.3223106

Zeng, K., Yu, J., Wang, R., Li, C., & Tao, D. (2017). Coupled Deep Autoencoder for Single Image Super-Resolution. IEEE Transactions on Cybernetics, 47(1), 27–37. 10.1109/TCYB.2015.2501373

Zhang, K., Liang, J., Gool, L. V., & Timofte, R. (2021). Designing a Practical Degradation Model for Deep Blind Image Super-Resolution (No. 2103.14006). arXiv. 10.48550/arXiv.2103.14006

Zhao, K., Pang, K., Hung, A. L. Y., Zheng, H., Yan, R., & Sung, K. (2025). MRI Super-Resolution With Partial Diffusion Models. IEEE Transactions on Medical Imaging, 44(3), 1194–1205. 10.1109/TMI.2024.3483109

Zhu, H., Li, Y., Ibrahim, J. G., Shi, X., An, H., Chen, Y., Gao, W., Lin, W., Rowe, D. B., & Peterson, B. S. (2009). Regression Models for Identifying Noise Sources in Magnetic Resonance Images. Journal of the American Statistical Association, 104(486), 623–637. 10.1198/jasa.2009.0029

